# PDE9 Inhibition Activates PPARα to Stimulate Mitochondrial Fat Metabolism and Reduce Cardiometabolic Syndrome

**DOI:** 10.1101/2021.02.02.429442

**Authors:** Sumita Mishra, Virginia S. Hahn, Nandhini Sadagopan, Brittany Dunkerly-Ering, Susana Rodriguez, Dylan C. Sarver, Ryan P. Ceddia, Sean Murphy, Hildur Knutsdottir, Vivek Jani, Deepthi Ashoke, Christian U. Oeing, Brian O’Rourke, Kavita Sharma, Jon Gangoiti, Dorothy D. Sears, G. William Wong, Sheila Collins, David A. Kass

## Abstract

Central obesity with cardiometabolic syndrome (CMS) is a major global contributor to human disease, and effective therapies are needed. Here, we show inhibiting cyclic-GMP selective phosphodiesterase-9A (PDE9-I) suppresses established diet-induced obesity and CMS in ovariectomized female and male mice. PDE9-I reduces abdominal, hepatic, and myocardial fat accumulation, stimulates mitochondrial activity in brown and white fat, and improves CMS, without altering activity or food intake. PDE9 localizes to mitochondria, and its inhibition stimulates lipolysis and mitochondrial respiration coupled to PPARα-dependent gene regulation. PPARα upregulation is required for PDE9-I metabolic efficacy and is absent in non-ovariectomized females that also display no metabolic benefits from PDE9-I. The latter is compatible with estrogen receptor-α altering PPARα chromatin binding identified by ChIPSeq. In humans with heart failure and preserved ejection fraction, myocardial expression of *PPARA* and its regulated genes is reduced versus control. These findings support testing PDE9-I to treat obesity/CMS in men and postmenopausal women.

**Summary:** Oral inhibition of phosphodiesterase type 9 stimulates mitochondrial fat metabolism and lipolysis, reducing central obesity without changing appetite

## INTRODUCTION

Nearly one out of every five humans around the globe is obese*(1)*, and in the United States this proportion exceeds 40% *(2)*. The pandemic has already had a major impact on global health as obesity increases the risks for diabetes, dyslipidemia, non-alcoholic fatty liver disease, inflammatory syndromes, heart failure, and hypertension, all components of cardiometabolic syndrome (CMS)*(1, 3, 4)*. Human studies have shown abdominal (visceral) obesity is most pathogenic*(5)*, whereas metabolically active adipose tissue known as brown fat appears to be protective*(6)*. Obesity-related disorders exhibit a sexual dimorphism as premenopausal women are relatively protected largely due to estrogen*(7, 8)*, yet after menopause when visceral fat increases, so do the risks for developing CMS*(9, 10)*. Obesity is now a defining feature of a common syndrome known as heart failure with preserved ejection fraction or HFpEF*(9, 11)*, which has high morbidity and little effective therapy *(10, 12)*. Shortly after the international spread of a novel Sars-COV2 coronavirus (COVID-19) in early 2020, obesity quickly emerged as an independent risk factor for worse disease and mortality with many of the mechanisms proposed being identical to those highlighted in CMS*(13)*. Despite its broad impact on human disease, effective obesity treatments remain limited. While diet and exercise are important, their impact is often limited in severely obese individuals given their low basal metabolic rates and exertional incapacity. Diabetes therapies such as glucagon-like peptide-1 agonists and sodium-glucose co-transporter-2 antagonists are being tested, but methods that stimulate fat catabolism and target visceral fat remain lacking.

An intrinsic lipolytic pathway is one coupled to cyclic GMP-protein kinase G (PKG) signaling stimulated by natriuretic peptides (NP) synthesized by the heart, or by nitric oxide. PKG is the primary kinase effector of cGMP and it phosphorylates hormone sensitive lipase (HSL) and perilipin, stimulates mitochondrial biogenesis and oxidative activity, and improves insulin signaling to suppress diet-induced obesity *(14, 15)*. Cyclic GMP also results in increased expression and activity of peroxisome proliferator activator-receptor alpha (PPARα)*(16)*, a master controller of mitochondrial fatty acid metabolism/oxidation and inducer of thermogenic (browning of) adipose tissue. PKG can be exogenously activated either by stimulating cGMP synthesis or suppressing its hydrolysis via specific phosphodiesterases (PDEs), and while there is shared downstream signaling, there are also many differences due to intra-cellular compartmentation and cell/organ specificity*(17)*. Stimulatory approaches include exogenous natural or synthetic NPs *(18)*, nitrates/nitrites*(19)*, and direct soluble guanylate cyclase stimulators*(20)*. However, each has limitations for chronic obesity therapy as they act rapidly, are short-lived and can potently lower blood pressure from vasodilation. NPs must also be injected.

PKG can also be stimulated by blocking cGMP-selective PDEs. The best-known example is PDE5 inhibition currently used to treat erectile dysfunction and pulmonary hypertension. PDE5-I also ameliorates cardiac pressure-overload*(21)* and is also reported to stimulate fat metabolism and induce metabolic activation of adipose tissue*(22)*. However, studies show this PDE primarily modulates NO-dependent cGMP, and when this signaling is diminished as occurs in females lacking estrogen, the impact of PDE5 inhibition is also very reduced*(23)*. Given most women with obesity-CMS are postmenopausal, this poses a major limitation. PDE9 is the other highly cGMP-selective PDE, and its inhibitors improve hearts subjected to pathological pressure-overload*(24, 25)*, hemodynamics and renal function in dilated heart failure*(26)*, and diastolic dysfunction*(27)*. While not yet approved in humans, several PDE9 inhibitors are currently in clinical trials for neurocognitive disease and heart failure. Unlike PDE5, PDE9 regulates NO-independent cGMP*(24, 28)*. PDE9 is expressed in adipose tissue, liver, and hearts of mammals including humans *(24, 29)*, though less so in systemic arteries, and its inhibition has little effect on blood pressure in animals*(24, 27)* and humans. However, the role of PDE9 in regulating fat metabolism and the potential for selective inhibitors to counter obesity and/or CMS are unknown.

Accordingly, the present study tested the capacity of a highly selective PDE9-I (PF-04447943) previously tested in humans with Alzheimer’s Disease*(30)* to counter diet-induced morbid obesity and CMS. We particular studied effects in females lacking estrogen and males, as these are the main groups impacted by this syndrome. In both, we find PDE9 inhibition reduces obesity and lipid accumulation in multiple tissues, the enzyme localizes to mitochondria and its inhibition stimulates mitochondrial respiration and fatty acid oxidation in conjunction with PPARα stimulation. We also find a sexual dimorphism, as obese intact females lack PDE9-I associated increases in PPARα-regulated genes and have no change in obesity-CMS.

## RESULTS

### Model of Combined Severe Diet-induced Obesity with Cardio-Metabolic Syndrome

To test the impact of PDE9-I on obesity-CMS, we subjected C56BL6/N mice to diet-induced obesity (DIO) for 6 months, with mild mechanical cardiac pressure-stress superimposed starting at month 5 and a week later, randomizing them to either PDE9-I or placebo (vehicle) treatment (Figure 1a). In OVX females, DIO resulted in substantial weight gain (100-200% over baseline) similar to that found in males (Figure S1a), and reflecting severe obesity (equivalent to a body mass index (BMI) >35 kg/m^2^ in humans). These mice had marked abdominal fat accumulation (Figure S1b), glucose intolerance (Figure S1c), and hepatic steatosis (Figure S1d).

**Figure 1.**
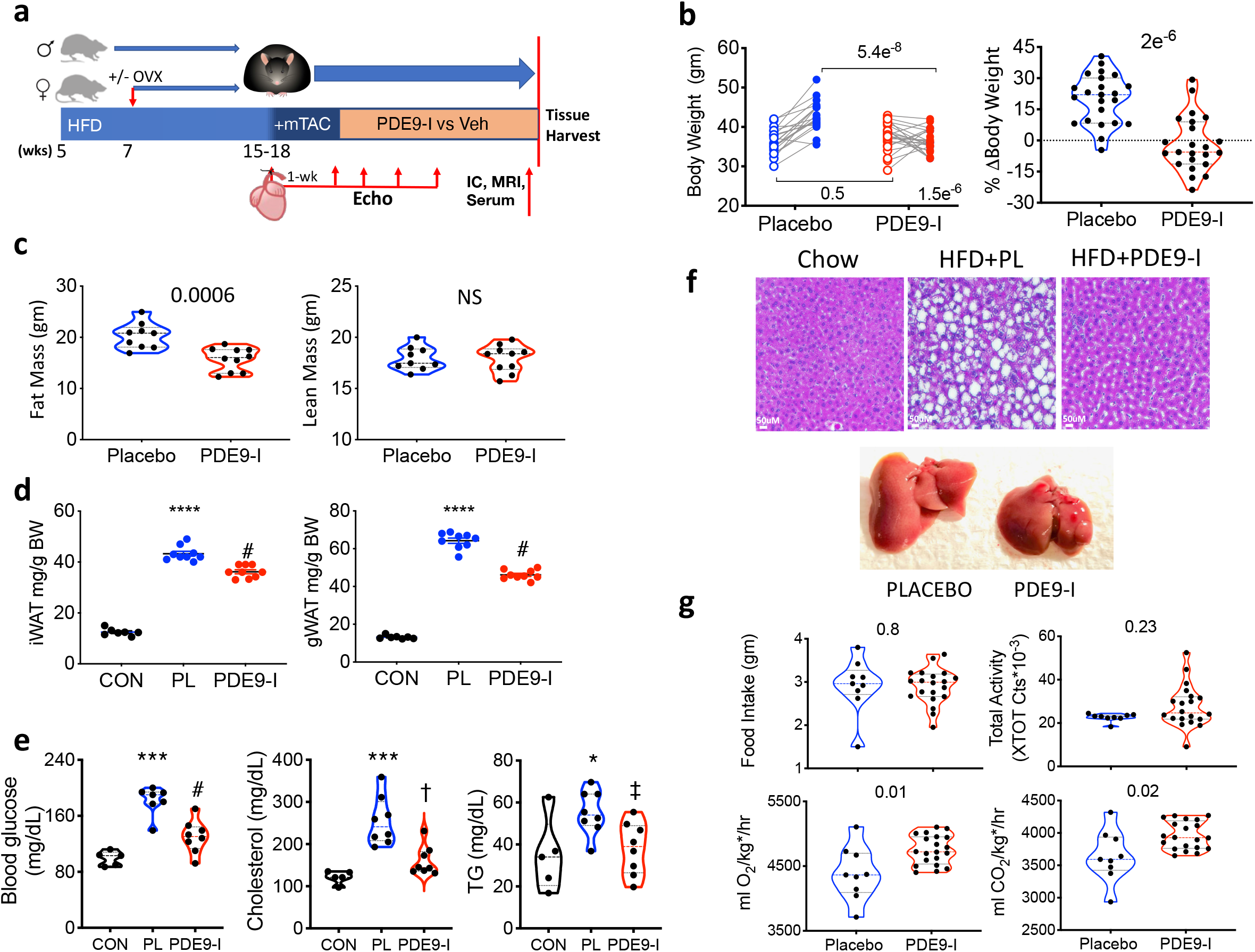
PDE9-inhibition suppresses diet-induced obesity and metabolic defects in ovariectomized females with cardiometabolic syndrome. **a)** Scheme of the protocol used to generate cardiometabolic syndrome: high-fat diet. (HFD), +/− ovariectomy (OVX) in females, subsequent mild aortic constriction (mTAC) to increase cardiac load, and lastly placebo-controlled p.o. PDE9-I drug trial (PF-04447943). IC – indirect calorimetry, MRI – magnetic resonance imaging for body fat-lean tissue, EC – echocardiogram, Veh-vehicle. **b) *Left*** Body mass in OVX mice before and after 8-wks of placebo (n=19) or PDE9-I (n=23) treatment (2WANOVA, with Sidaks multiple comparison). ***Right***: Percent change in body mass between time points. (Mann Whitney, MW test). **c)** Magnetic resonance imaging-derived total body fat and lean mass for two treatment groups in OVX mice. (p-values, MW). **d)** Inguinal (iWAT) and gonadal (gWAT) white adipose tissue weight in OVX females treated with placebo or PDE9-I (n=7-9/group, KW, Dunns multiple comparisons test (DMCT); **** p=0.00001 vs normal diet control (CON), # p=0.028 vs placebo). **e)** Serum fasting blood glucose and lipids in OVX mice on control versus HFD +/− PDE9-I; (ANOVA, Holm Sidak (HS) comparisons test, *** p=0.0004, * p=0.03 vs CON; # p=0.03, †p=0.037, ‡ p=0.047 vs Placebo (PL)). **f) Upper:** Representative liver histology in OVX with standard diet (Chow) or HFD with placebo vs PDE9-I. Marked steatosis was seen with placebo treatment that reverted to near normal by PDE9-I. (replicated n=5/group). **Lower** Examples of whole liver showing a marked reduction of liver mass by PDE9-I. Group results in Figure S2a. **g)** Food intake, activity, and whole-body respiration in OVX mice +/− PDE9-I; (MW). O2 and CO2 consumption/production rates are normalized to mass kg* = (lean mass + fat mass*0.2) as described*(31)*.

### PDE9-I lowers obesity and improves CMS in OVX-female and male mice

Using this obesity-CMS model, we tested the impact of PDE9-I on total body fat and lean mass and metabolic dysregulation in OVX females. Mice receiving placebo continued to gain weight (median 22%) during the 8-week trial, whereas PDE9-I treated mice exhibited a net −5.6% weight decline (p=2 •10^−6^, Fig 1b). Of the 23 mice treated with PDE9-I, 14 (61%) experienced a 2% or greater decline in body weight versus 1/20 (5%) in placebo (p<0.0001). MR body composition analysis revealed that PDE9-I treated mice had less total fat mass (primarily abdominal) and unaltered lean mass (Figure 1c), and less inguinal (subcutaneous) and gonadal (visceral) fat pad mass (Figure 1d). Metabolic defects including elevated fasting blood glucose, triglycerides, and cholesterol that were present in the placebo group were all reduced with PDE9-I (Figure 1e). Hepatic steatosis and liver mass were greater in OVX placebo-treated mice but were at near-normal levels with PDE9-I in both OVX (Figure 1f, Figure S2a, S2b) and male (Figure S2c) mice. Importantly, PDE9-I did not significantly change food intake or activity level over placebo control. However, total body O2 consumption and CO2 production were significantly greater in PDE9-I treated mice (Figure 1g) without a change in core temperature (Figure S2d).

Age-matched males subjected to the same obesity/CMS model and then treatment protocol also displayed lower total body weight primarily due to reduced fat mass (Figure S3a,S3b) and less dyslipidemia (Figure S3c) when provided PDE9-I. PDE9-I also did not significantly alter food intake or activity but increased both body O2 consumption and CO2 production in males (Figure S3d) as it did in OVX females.

### PDE9-I improves cardiac function, and blunts hypertrophy/fibrosis gene expression and augments myocardial PPARα-signaling relative to placebo in obese-CMS mice

Prior studies have shown PDE9 counters cardiac hypertrophy and depresses pro-fibrotic signaling cascades in non-obese males subjected to pressure-load stress, and this efficacy is independent of NO-signaling*(24)*. Whereas PDE5A-I does not augment myocardial cGMP in OVX females*(23, 32)*, here we show PDE9-I does (Figure S4a), and this suggested it might also counter cardiac stress in our model. PDE9-I improved ejection fraction over placebo in OVX mice (Figure 2a, p=0.001 for drug/group interaction) and attenuated a significant rise in LV mass with placebo (p=0.005 for interaction). Similar effects were observed in males (Figure S4b). Diastolic function was impaired in the OVX placebo group and this improved to lean control levels with PDE9-I (Figure 2b). PDE9-I also lowered pro-hypertrophic and pro-fibrotic related gene expression compared with placebo in OVX females and males (Figure 2c, Figure S4c).

**Figure 2.**
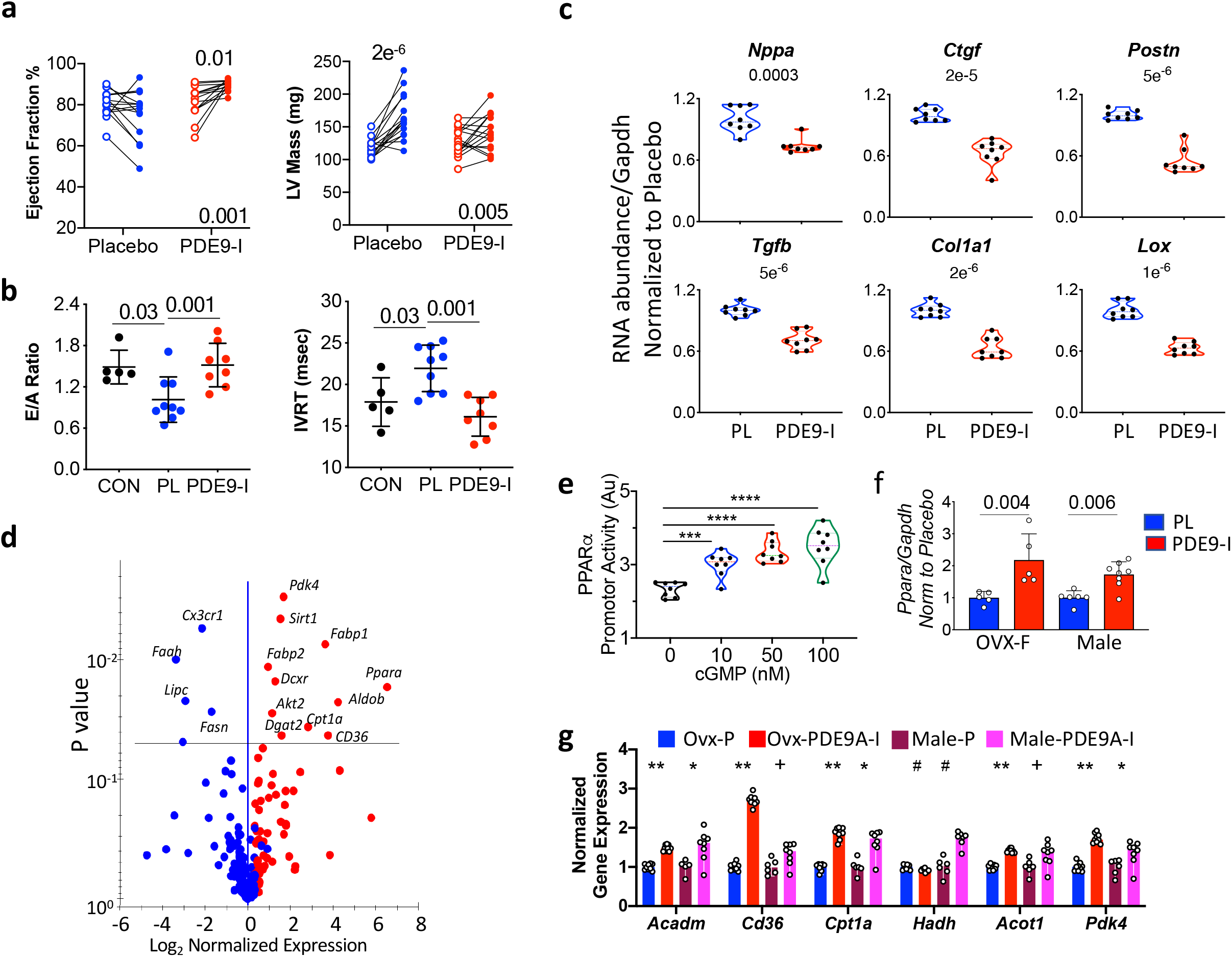
PDE9-I improves cardiac function, suppress pathological hypertrophic gene expression, and stimulates PPARα signaling in OVX myocardium. **a)** Left ventricular ejection fraction and left-ventricular (LV) mass in placebo vs PDE9-I treated OVX mice, with paired data at onset of end of 8 wk treatment period (n=15,16 for placebo; PDE9-I; RMANOVA, p-value lower right for treatment x time interaction, above for post-hoc Tukey test within group comparisons). **b)** Diastolic function assessed by mitral filling ratio (early/atrial; E/A) and isovolumic relaxation time (IVRT) in normal diet controls (CON), and obese-CMS OVX mice treated with placebo (PL) or PDE9-I. n=5-9/group, Kruskal Wallis, Dunns post-hoc comparison. **c)** mRNA abundance normalized to *Gapdh* for A-type natriuretic peptide *(Nppa)*, transforming growth factor beta *(Tgfb)*, collagen type 1a1 *(Col1a1)*, lysyl oxidase *(lox)*, connective tissue growth factor *(ctgf)* and periostin *(postn)* in OVX myocardium, (8/group, Mann Whitney t-test). **d)** Volcano plot of differential gene expression in OVX myocardium between placebo and PDE9-I treatment, using lipid and carbohydrate metabolism PCR array. Benjamini-Hochberg adjusted p-value versus log-2 fold change is shown (n=10/group). **e)** Activation of PPARα promoter by cGMP in Hep G2 cells; (n=8/group, ANOVA, Holm Sidak comparisons test; *** p=0.007; **** p≤2e^-6^). **f)** *PPARα/Gapdh* gene expression in brown adipose tissue (BAT) from obese-CMS OVX and males treated with placebo versus PDE9-I. (2-step Benjamini, Krieger, Yekutieli multiple comparisons KW test (2-St-BKY-MC-KW) test, q-values shown) **g)** PPARα-regulated fatty-acid metabolism genes expressed in myocardium in OVX and male mice with placebo (P; n=9,6) or PDE9-I (n=9,8); 2-St-BKY-MC-KW. Q-values: ** p=0.00005; # p=0.008; + p=0.03; * p=0.02 vs placebo).

Increased dietary nitrate is reported to enhance fatty acid (FA) oxidation by increasing the expression of PPARα and associated downstream signaling to augment mitochondrial respiration and fat metabolism*(16)*. To see if PDE9-I triggers a similar cascade, we first performed a targeted PCR-array analysis of heart tissue finding an abundance of transcripts for PPARα and many of its target FA-regulating genes increased (Figure 2d). Cyclic GMP increased PPARα promotor activity in Hep G2 (liver) cells in a dose-dependent manner (Figure 2e). Concordantly, PPARα mRNA increased in the myocardium of OVX and male mice treated with PDE9-I (Figure 2f). Myocardial PPARα activity was assessed by mRNA abundance of regulated genes involved with FA metabolism. This increased in both OVX and male mice (Figure 2g). Metabolite profiling for myocardial acylcarnitines revealed greater levels primarily of long-chain FA in both OVX and male mice receiving placebo as compared to controls. These levels were less in both PDE9-I treated groups (Figure S4d). Collectively these results show PDE9-I ameliorates cardiac dysfunction and remodeling, reduces pro-fibrotic signaling, and enhances FA metabolism in concert with increased PPARα signaling.

### PDE9-I induces adipose tissue browning and enhances fatty acid catabolism

Activation of FA oxidation by PPARα manifests in brown (BAT) and white (WAT) adipose tissue by increased mitochondria biogenesis and fat oxidation. Figure 3a shows example histology of BAT from a non-obese control and obese OVX and male mice treated with placebo or PDE9-I. BAT from lean mice has a darker color due to small lipid droplets and increased mitochondria density (left panel). With obesity-CMS, BAT adipocytes in placebo-treated OVX and males were enlarged with more lipid storage (pale color). This appearance was much closer to normal with PDE9-I. BAT from PDE9-I treated mice also had increased mRNA abundance of multiple genes regulating FA metabolism and mitochondrial respiration (Figure 3b) accompanied by a marked decline in acylcarnitines to levels found in lean controls (Figure 3c). Genes regulating mitochondrial biogenesis increased consistent with fat browning (Figure 3d). PDE9-I treatment also enhanced expression of FA metabolism regulating genes in WAT (Volcano plot, Figure 3e) and in genes controlling mitochondrial respiration and uncoupling (Figure 3f). These results show PDE9-I activates adipocyte catabolic and thermogenic programs reversing abnormalities due to severe obesity.

**Figure 3.**
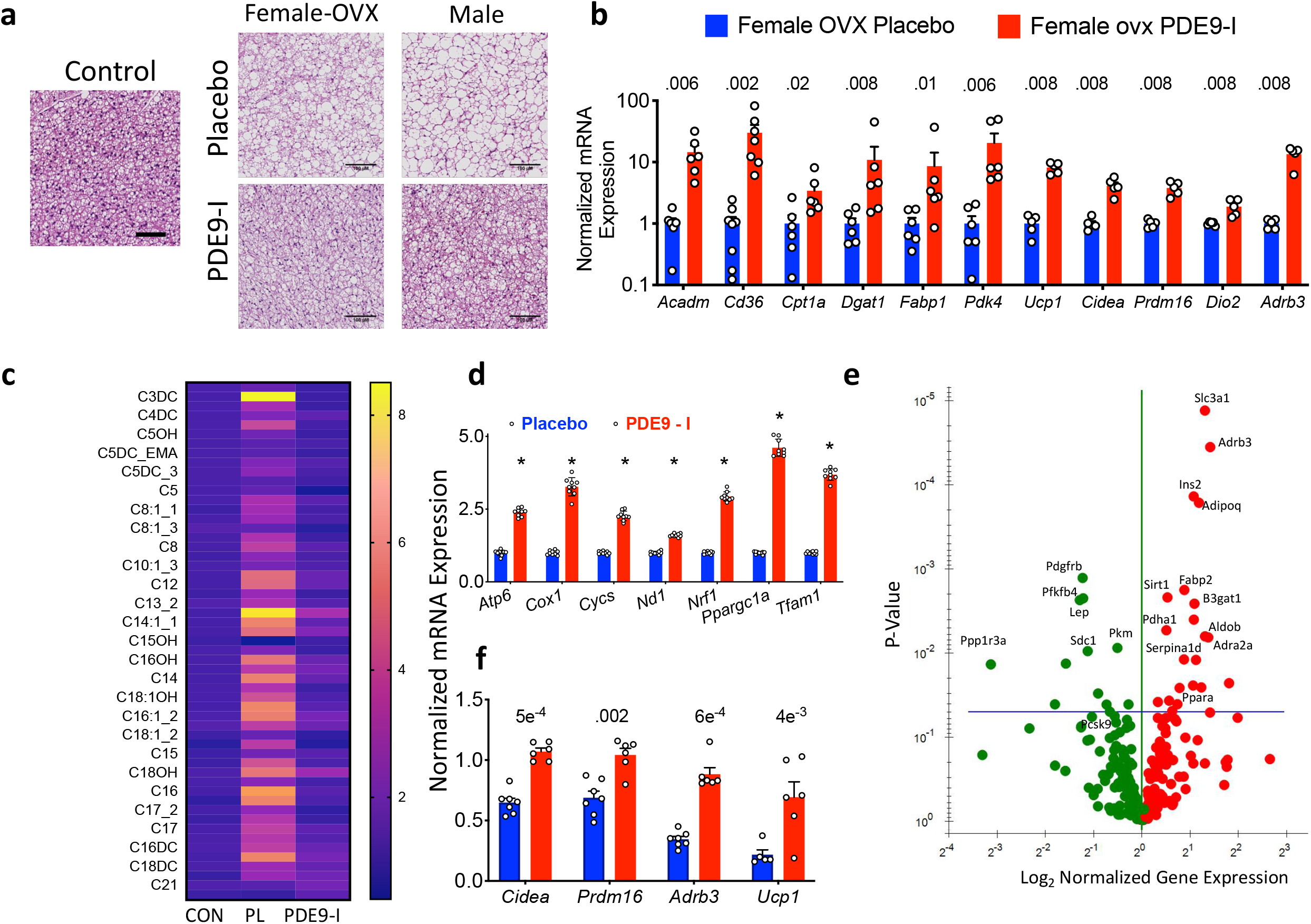
PDE9-I reduces multi-organ lipid accumulation by enhancing lipolysis and mitochondrial respiration. **a)** Representative histology of brown fat in a normal control (left), and obese-CMS OVX mice after placebo or PDE9-I treatment (replicated n=7/group; scale bar 100 μm). **b)** BAT mRNA abundance for fat metabolism and mitochondrial respiration genes in OVX mice treated with placebo or PDE9-I. 2-St-BKY-MC-KW q-values shown. **c)** Metabolomic analysis of acylcarnitines in BAT from OVX obese-CMS mice with placebo (PL) or PDE9-I treatment. N=5 per group, 2-St-BKY-MC with Mann Whitney test, all p values <0.008). **d)** mRNA abundance of mitochondrial biogenesis and oxidative activity genes in BAT from OVX with PL or PDE9-I. n=9/group, * q<1e^-11^ between groups, same analysis as for panel **b. e)** Volcano plot of differential gene expression for lipid metabolism PCR array in WAT tissues from OVX model treated with PDE9-I versus placebo. Benjamini-Hochberg adjusted p-value versus log-2-fold change is shown (n=8/group). **f)** mRNA abundance of mitochondrial oxidative genes in WAT from OVX treated with placebo or PDE9-I. (n=5-7/group, same stats as in S3b).

### PDE9-I directly stimulates fat lipolysis and increases mitochondrial respiration

The decline in adipose tissue and hepatic fat infiltration by PDE9-I suggests it can stimulate lipolysis in multiple cell types. In cardiomyocytes pretreated with vehicle or PDE9-I and then fed FFA for 48 hours, PDE9-I resulted in reduced intra-cellular lipid accumulation (Figure 4a, 4b). Using a glycerol release assay, we found PDE9-I alone induces lipolysis in differentiated adipocytes (Figure 4c). Dose-dependent increases in glycerol release by PDE9-I were also observed in liver Hep G2 cells and cardiomyocytes, but this required ANP co-stimulation. Enhanced lipolysis by PDE9-I was blocked by co-inhibition of PPARα (Figure 4d). In addition, the PDE9-I effect was negated in cells pre-incubated with *PDE9* siRNA confirming selectivity of the pharmacological inhibitor.

**Figure 4.**
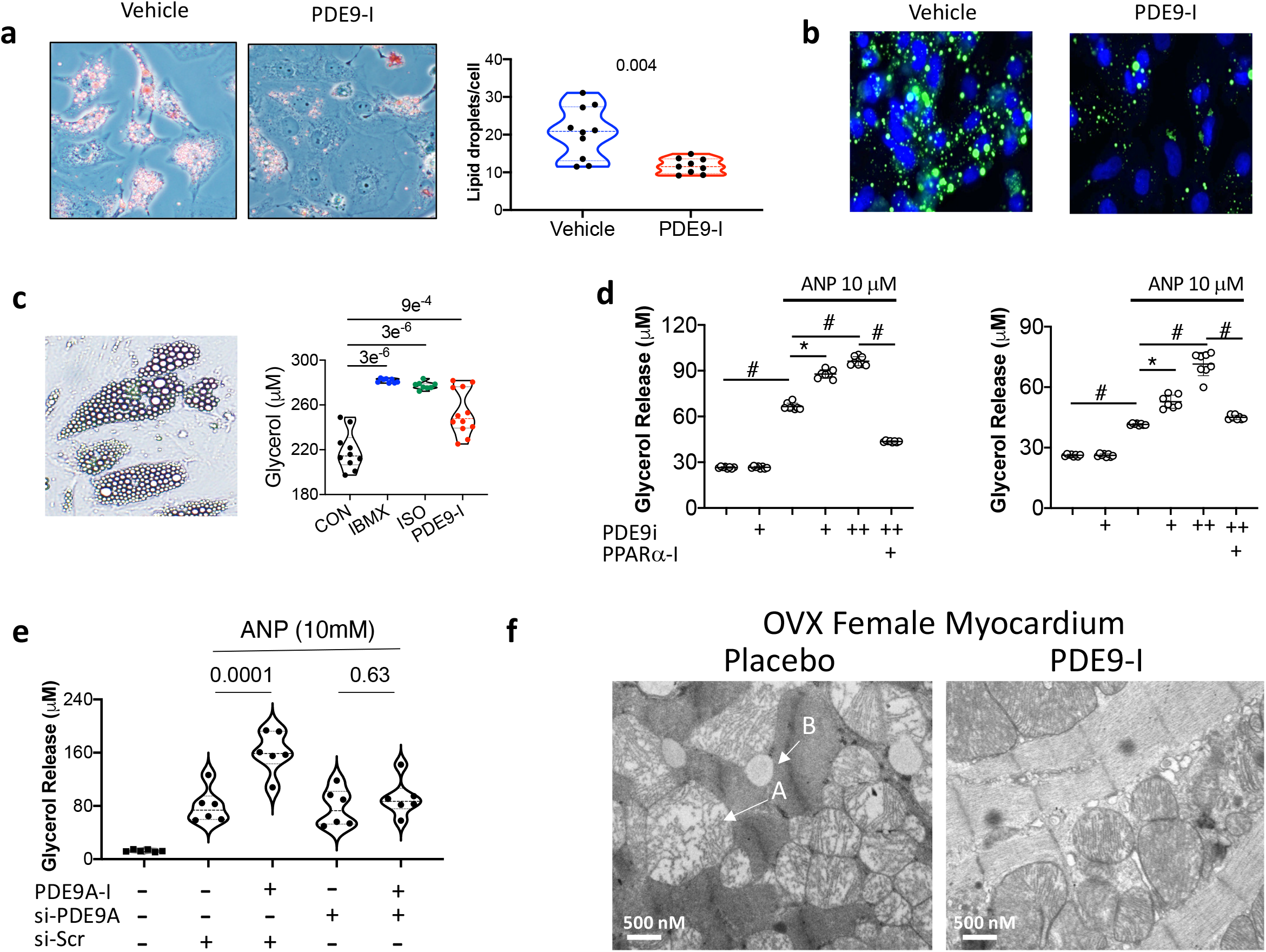
PDE9-I increases lipolysis in fat, liver and myocardium. **a)** Phase contrast microscopy with Oil Red-O in cardiomyocytes fed a lipid mixture for 48 hrs and co-treated with vehicle or PDE9-I (replicated 4x), and summary for lipid droplets/myocyte on right (n=10/group, MW). **b)** Same protocol with myocytes loaded with 5 μM BODIPY 493/503 labeling lipid droplets green and nuclei (DAPI) blue; (x3 replicates). **c)** Induction of lipolysis (glycerol released) in 3T3 L1 derived adipocytes incubated with IBMX (broad PDE inhibitor), isoproterenol (ISO), or PDE9-I. (n=10,8,8,12 respectively; Welch ANOVA, DMCT). **d)** Lipolysis with PDE9-I +/− ANP stimulation in HepG2 (left) and myocytes (right). PDE9-I (+ 2.5 μM, ++ 5 μM) stimulated lipolysis in the presence of ANP, and this was suppressed by concomitant PPARα inhibition. # p=0.0006; * p=0.0012, KW unpaired-test with Bonferoni correction for 4 comparisons. **e)** PDE9-I augmentation of ANP stimulated lipolysis is prevented in HepG2 cells expressing siRNA-*PDE9* vs siRNA-*scrambled*. (n=6/group; 2WANOVA with Tukey comparisons test). **f)** EM images of LV myocardium from OVX ob/CMS mice treated with placebo or PDE9-I. Disruption of normal structure with separation of cristae and reduced density (arrow-A) and lipid accumulation (arrow-B) are shown with placebo, but much less evident with drug treatment.

Diet induced obesity increases triglyceride storage in multiple tissues causing lipotoxicity that can alter mitochondrial architecture and function, inducing swelling with reduced cristae density *(33, 34)*. EM revealed such mitochondrial changes as well as myocardial lipid accumulation in both OVX and male mice treated with placebo, and both features improved with PDE9-I (Figure 4f, Figure S5a, S5b). Together, these data show PDE9-I stimulates lipolysis particularly with NP background stimulation, reducing mitochondrial lipid accumulation and morphological disruption.

### PDE9 localizes to mitochondria and stimulates fat oxidation

We previously reported PDE9 co-localizes with the sarcoplasmic reticular (SR) ATPase at T-tubules in cardiac myocytes in contrast to PDE5A which localizes to the Z-disk*(24)*. The proximity of T-tubular membranes and mitochondria in the dyadic cleft raises the possibility that PDE9 also localizes with the latter. To test this, *PDE9*-*Flag* was expressed in cardiomyocytes and cells fractionated into mitochondrial and cytosolic components. We used Flag-Ab as native PDE9 antibody detection is poor with existing reagents. PDE9-FLAG was enriched in the mitochondrial fraction (Figure 5a) even after normalization for total protein. By contrast, *Pde5a-Flag* was primarily found in the cytosol (Figure S6a). Secondly, mitochondria were stained with Mitotraker Red which colocalized with a *PDE9-Gfp* imaged with confocal fluorescence microscopy (Figure 5b). Lastly, immuno-gold electron microscopy of cardiomyocytes expressing GPF-PDE9 or GFP revealed a diffuse cytosolic distribution for GFP but primarily mitochondrial localization for PDE9-GFP (Figure 5c). Together these results show PDE9 can specifically localize at mitochondria.

**Figure 5.**
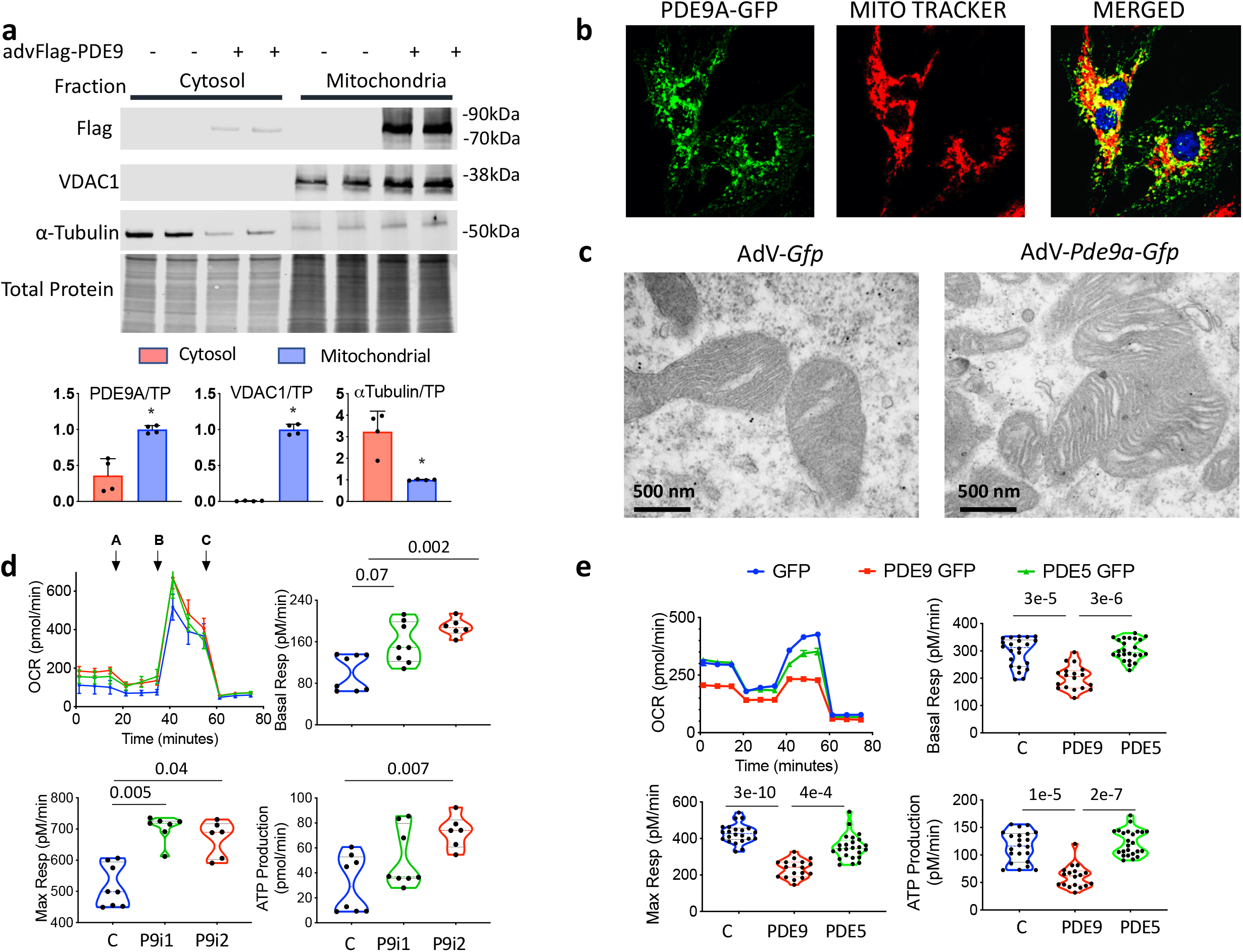
PDE9 localizes to mitochondria and its inhibition augments fat oxidation. **a)** Cardiac myocytes expressing PDE9-FLAG show higher levels in mitochondrial versus cytosolic fraction. Summary results n=4, *-p=0.028 by MW. **b)** Co-localization of PDE9-GFP with Mitotracker-Red by confocal fluorescence imaging (repeated x3). **c)** Example - gold labeled PDE9-GFP and GFP imaged by EM. Dark black dots identify the proteins. GFP is diffusely distributed whereas PDE9-GFP is primarily found in mitochondria. (repeated x12/group). Lower magnification images shown in Figure S6b. **d)** Oxygen consumption rate (OCR) in myocytes pretreated with vehicle (Veh, n=8) or PDE9-I (P9i-1 2.5μM (n=8), 5μM (n=6) x 24 hrs and then fed fatty acids as primary substrate x 24 hrs. A:oligomycin; B:FCCP; C:rotenone + antimycin (n=8,6; KW, DMCT p values displayed) **e)** OCR in myocytes pre-infected with AdV expressing *PDE9-Gfp, Pde5a-Gfp*, or *Gfp* alone for 48 hrs. (n=18,24, 22 respectively; KW, DMCT p-values displayed).

To test the impact of PDE9 loss-of-function on mitochondrial FA oxidation, cardiomyocytes were pre-treated with PDE9-I or vehicle and then incubated with palmitate the primary fuel. Basal and maximal oxygen consumption and ATP production were higher in PDE9-I treated cells (Figure 5d). Gain-of-function was assessed by pre-incubating myocytes with Adenovirus expressing either *Gfp-Pde9a* or *GFP-Pde5a*. PDE9 protein expression lowered basal and maximal respiration and ATP production, whereas these were unchanged with PDE5A overexpression (Figure 5e). Together, these data show PDE9 localizes to mitochondria, and that its expression inhibits respiration and ATP generation whereas its inhibition achieves the opposite.

### PPARα signaling is downregulated in Human HFpEF

Among the most prominent clinical syndromes combining morbid obesity and CMS is HFpEF, which now affects over half of all patients with heart failure*(11)*. Our RNAseq analysis of differentially expressed genes in myocardium from human HFpEF versus controls found the transcriptome is potently influenced by BMI, and identified patient subgroups with distinct signaling pathway and clinical characteristics*(35)*. Using this database, we now show mRNA abundance for *PPARA, PPARG*, and *PPARGC1A* is reduced by ∼40% (adjusted q-score, p<1e^-6^ for each, Figure 6a) in HFpEF versus controls. A volcano plot of genes associated with PPAR signaling (based on KEGG) shows those coupled to fatty acid uptake *(FABP5, FABP4, CD36)*, degradation *(LPL)*, and oxidation *(CPT1a, ACOX1/3, ACAA1, ACSL3/4)* are significantly reduced in HFpEF (Figure 6b). Unsupervised hierarchical cluster analysis using *PPARA* and *PDE9A* expression only identified two patient groups (Figure 6c), both similarly obese (Table S1). *PDE9A* was similarly and directly correlated with *PPARA* abundance in each group, but Group 1 had significantly lower *PPARA* expression for any given *PDE9A* level (P=7e^-8^ for offset by ANCOVA). Group 1 had fewer patients with diabetes, their left ventricles were smaller with less hypertrophy, and plasma natriuretic peptide levels lower as compared to Group 2. Interestingly, Group 1 overlaps with a HFpEF subgroup we had previously identified using agnostic non-negative matrix factorization and the broad transcriptome *(35)*, that contained mostly females with smaller hearts and pro-inflammatory signaling (Table S1). Group 2 overlaps with a second sub-group whose hearts display expression profiles closest to that for HF with low EF *(35)*. These results support the relevance of *PPARA* downregulation in HFpEF and reveal a subgroup that may be particularly responsive to enhancement by PDE9-I.

**Figure 6.**
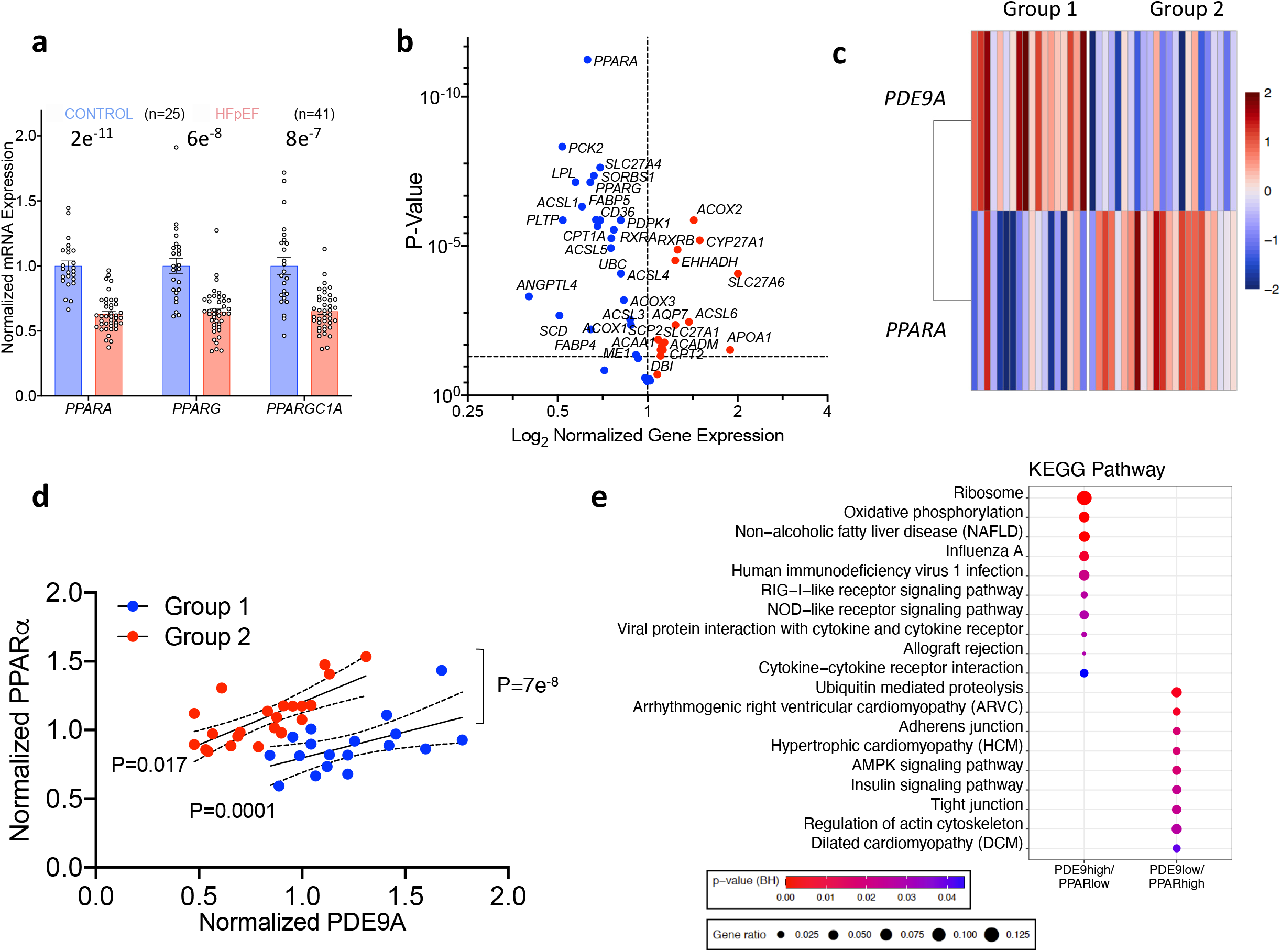
*PPARA* and related gene expression is reduced in human HFpEF myocardium directly relates to *PDE9A* expression, and defines subgroups. **a)** Expression of PPARa, PPARg, and PPARGC1A (PGC1a) in HFpEF (n=41) myocardium normalized to non-failing controls (n=24). (2S-BKY-MC-MW, q values shown). **b)** Volcano plot of differential mRNA expression for genes in the KEGG-PPAR pathway. Benjamini-Hochberg MCT – q-values on y-axis. **c)** Unsupervised, hierarchical heat map cluster analysis using *PPARA* and *PDE9* genes from the human HFpEF RNAseq data yielded two HFpEF groups with reciprocal relative expression of the two genes. **d)** Regression *PDE9* versus *PPARA* expression for sub-groups identified by the cluster map. Each group has a significant positive dependence with a similar slope but different offset (ANCOVA). **e)** KEGG pathway analysis from differentially expressed genes in the two HFpEF subgroups defined by relative *PPARA* versus *PDE9* expression levels.

### PPARα activation is required for PDE9-I to reduce obesity and CMS

Prior studies have shown activation of PPARα in models of DIO reduces fatty liver and can prevent development of obesity*(36, 37)*, while the PPARα KO mouse develops obesity gradually over many months*(38)*. To test if PPARα activity is required to drive the anti-obesity efficacy from PDE9-I, OVX females with obesity-CMS were treated with either PDE9-I alone or combined with a selective PPARα inhibitor (GW6471, daily i.p.). We chose this approach versus using a PPARα KO as we could generate the identical obesity-CMS condition prior to interfering with PPARα stimulation. GW6471 co-treatment reduced mRNA abundance of PPARα regulated genes by about 50%, supporting on-target effects (Figure 7a). As before, mice receiving only PDE9-I had less total body and fat mass but no significant change in lean body mass and increased VO2 and VCO2 versus placebo, and co-inhibition of PPARα prevented these changes (Figure 7b). Total food intake and daily activity did not significantly differ between the groups (Figure S7a). Compared with placebo, reduced iWAT and gWAT mass (Figure 7c) and improved cardiac fractional shortening (Figure 7d) with PDE9-I were largely abrogated by concomitant PPARα blockade. Thus, the *in vivo* efficacy of PDE9-I on obesity-CMS required PPARα activation.

**Figure 7.**
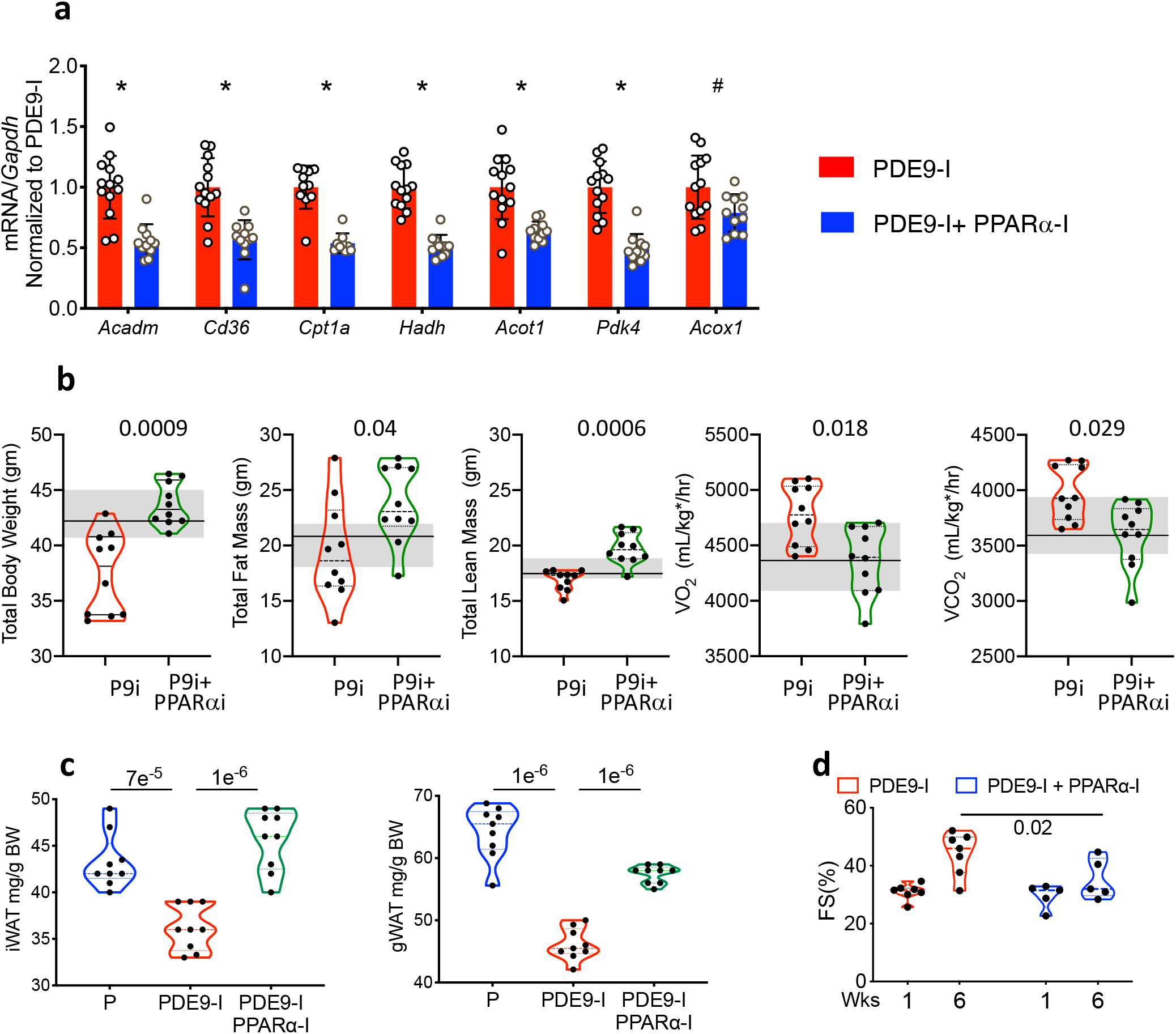
Inhibition of PPARa blocks beneficial effects of PDE9-I in OVX model. **a)** Quantitative PCR of PPARa-associated genes in myocardial tissue from OVX ob/CMS mice treated with PDE9-I ± PPARa-inhibition. 2S-BKY-MC-MW, q values * - p≤5e^-5^; # p=0.007. **b)** Effect of PPARα co-inhibition on PDE9-I induced reduction of total body, fat, and lean mass, and increases in VO2 and CO2 in OVX ob/CMS mice. Results for placebo treated OVX mice (derived from data in Figure 1) are plot with median (dark line) and 75%-25% confidence intervals (shaded). The addition of a PPARα-I significantly reversed PDE9-I mediated responses in these parameters, returning their values to those with placebo for all by lean mass that rose by 5%. (n=10/group, p-values displayed from KW (includes all 3 groups) and DMCT) **c)** iWAT and gWAT weight in OVX mice PDE9-I ± PPARα-I versus placebo; n=9/group, ANOVA, p-values from TMCT). **d)** Fractional shortening (%FS) from same experiment; (n=7,5, 2WANOVA, p-value from Sidak’s comparisons test).

### PDE9-I is ineffective in non-OVX females with ERα shifting PPARα DNA binding away from FAO regulating genes

Endogenous estrogen has been found to suppress PPARα-stimulated gene expression and corresponding regulation of FA oxidation in tissue and in vivo obesity models*(39)*. Consistent with this, the PPARα activator fenofibrate effectively reduced DIO in males and OVX females but non-OVX*(37)*. These results combined with the current data showing that PPARα inhibition prevents PDE9-I from reducing obesity, led us to hypothesize a similar sex dimorphism may apply to effects from PDE9-I. This was tested in a cohort of non-OVX female mice subjected to the identical HFD-mTAC protocol. As shown in Figure 8, PDE9-I had negligible impact on total body, fat, or lean mass (Figure 8a), BAT adipocyte size or browning (Figure 8b), serum lipids (Figure S7b), iWAT or gWAT mass (Figure S7c), food intake, activity, and whole-body respiration (Figure S7d), or hepatic steatosis (Figure S7e) in non-OVX females.

**Figure 8.**
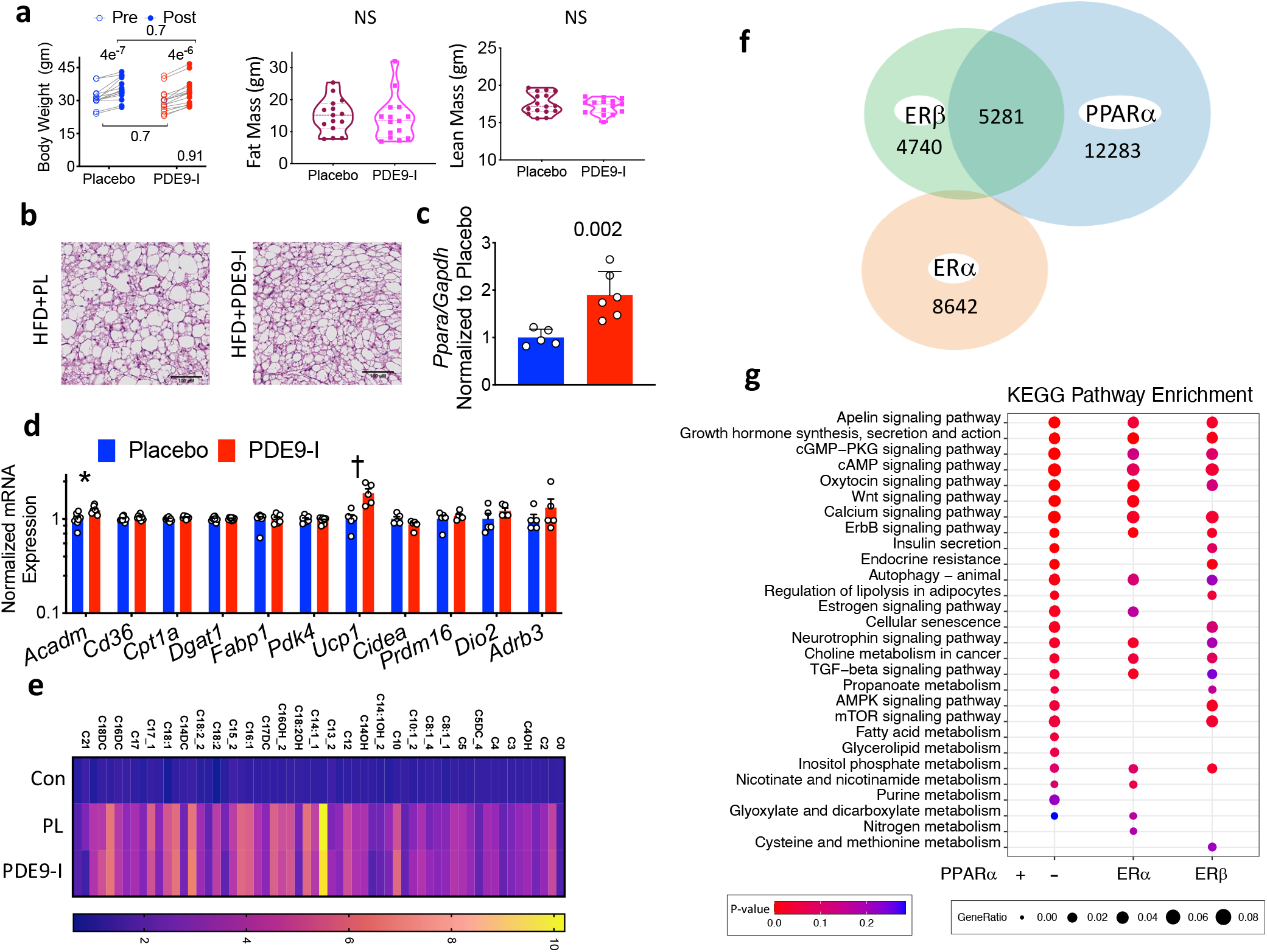
PDE9-I does not impact body weight, fat, or PPARa downstream signaling, associated with estrogen suppression of PPARα DNA binding to fat metabolism genes. **a)** Total body weight, fat, and lean mass are not significantly altered in non-OVX ob/CMS female mice treated with placebo versus PDE9-I. **b)** BAT histology shows enlarged adipocytes and reduced mitochondria density in the placebo group, that is not changed with PDE9-I (repeated x3) **c)** *PPARa* mRNA expression increases in non-OVX at levels similar to those for male and OVX mice. (MW). **d)** qPCR results for fatty acid metabolism and mitochondrial respiration genes in BAT from non-OVX model. (log-transformed, 2S-BKY-MC-MW, * q=0.03, † q=0.04, all others >0.5). **e)** Metabolomics of acylcarnitines in BAT from normal controls and ob/CMS non-OVX with placebo vs PDE9-I. (n=5/group, 2S-BKY-MC-MW; for placebo vs PDE9-I all q-values >0.5; for control chow (non-obese) versus placebo, all but one p≤0.001; (C22, p=0.006). **f)** Venn diagram of ChIP-seq identified PPARα binding sites in HepG2 cells with PPARα stimulation alone or combined with ERα or ERβ co-stimulation. **g)** KEGG pathway analysis from ChIP-seq identified genes with PPARα binding shows loss of fat metabolism related pathways by co-activation with ERα or ERβ.

Failure of PDE9-I to alter obesity-CMS in non-OVX females was not due to a lack of increased *Ppar*α mRNA abundance which rose similarly as in OVX females and males (Figure 8c, compare to Figure 2f). However, mRNA levels for genes controlling FA metabolism and mitochondrial oxidation/uncoupling downstream of PPARα were very little or not significantly changed (Figure 8d). Acyl-carnitine analysis of BAT also found no changes despite PDE9-I (Figure 8e).

These results indicate that sex hormones in intact females impeded the capacity of PDE9-I to increase PPARα-regulated FA catabolism. Since *Ppara* expression increased yet mRNA abundance for its regulated genes did not, we speculated that coincident estrogen receptor (ER) signaling alters PPARα DNA binding to impact the latter’s transcriptional function. To test this, ChIP-seq for PPARα binding was performed using Hep G2 cells transfected with either PPARα (+PPARα agonist) alone, or in combination with either ERα or ERβ transfection and respective agonists. We found ∼17,500 PPARα DNA binding sites when stimulated alone, but this number fell by nearly half and engaged mostly different sites if ERα was co-activated (Figure 8f). With ERβ co-activation, ∼1/3 of binding sites with PPARα alone were maintained, and 1/3 were altered; and there was also an overall reduction. Gene lists based on PPARα chromatin binding for the three conditions were subjected to KEGG pathway enrichment analysis, showing ERα co-activation particularly reduces PPARα binding to genes involved with FA metabolism (Figure 8g). This is consistent with the mRNA analysis (Figure 8c, d) and the lack obesity-CMS effects by PDE9-I in non-OVX mice.

## Discussion

This study supports the utility of a new treatment for severe obesity and associated cardiometabolic syndrome based on PDE9 inhibition. We show PDE9 localizes to mitochondria, that its suppression increases fat-metabolism and lipolysis, and reduces fat deposits in the abdomen, liver, and myocardium. Concomitant reduction of myocardial lipids particularly long-chain acylcarnitines is important as their accumulation is associated with cardiac and lipotoxicity with mitochondrial dysfunction *(34)*. The ability of PDE9-I to reduce fat accumulation in the liver and abdomen is also significant, as both depots are linked to worse CMS morbidity*(40)*. Cardiac pathobiology common with CMS, such as LV hypertrophy and pathological pro-fibrotic signaling were also ameliorated by PDE9-I. The link between PDE9-I and *Ppar*α upregulation involves promoter activation to increase expression of its many regulated genes controlling fat metabolism, mitochondrial respiration and biogenesis. The metabolic effects from PDE9-I are absent in non-OVX females despite increased PPARα mRNA, as the downstream gene activation does not occur. This is compatible with a shift of PPARα chromatin binding away from genes controlling fat-metabolism by the co-activation of ERα. Importantly, we also find depressed myocardial *PPARA* expression and associated signaling in obese humans with HFpEF. Given that PDE9 inhibitors appear safe and well tolerated in humans based on prior testing for other disorders, the current findings support their clinical assessment in men and postmenopausal women with obesity-CMS.

Studies first reporting lipolytic and anti-obesity effects from PKG stimulation identified several mechanisms, including phosphorylation and activation of hormone sensitive lipase, p38-MAP kinase (by unknown intermediate)*(41)* similar to the pathway used by PKA*(42)* and AMP activated kinase*(43)*. These are felt to converge on transcriptional regulators PPAR gamma co-activator 1α (PGC1α) and PPARα*(14, 15)* to control metabolism. PKG-dependent PPARα activation has been previously linked to mitochondrial protection against hypoxia-induced cardiac injury*(19)* and increased FA oxidation and catabolism in skeletal muscle*(16)*, though not with countering obesity. To our knowledge, none of these kinase or lipase effectors of PKG modulation have been shown to confer a sexual dimorphism on metabolic signaling. However, a sexual dimorphism has been reported for the impact of PPARα activation, as obese OVX females and males show lipolysis and reduced weight, whereas non-OVX do not *(37, 44, 45)*. A similar disparity in PDE9-I responses coupled to differential downstream PPARα signaling supports this factor as central to the metabolic and anti-obesity effects. Estrogen itself stimulates fat metabolism and mitochondrial oxidative respiration*(8)*, and its decline after menopause or OVX is associated with abdominal obesity, CMS, and reduced PPARα and PGC1α regulated signaling*(7)*. ER and PPARα have similar DNA binding motifs and protein partners such as RXR and coactivator PGC1α*(45)*, and prior studies have reported ERα protein interaction with PPARα and inhibition of latter’s expression and/or function in liver cancer*(46)* and in apolipoprotein regulation*(47)*. To our knowledge, the present analysis is the first to directly test this interaction with ChIPSeq. The marked shift in PPARα binding by ERα co-activation was striking. While many pathways remained engaged, those involved with fat metabolism declined. This suggests that pre-menopause, ER signaling may substitute for PPARα regulation, whereas post-menopause, PPARα becomes prominent so its activation by agonists or PDE9-I counters obesity. The mechanisms governing the chromatin interactions between PPARα and ERs, and their testing in other cell types, remain to be explored.

While our results show cGMP-PKG activates PPARα expression and gene regulation, other mechanisms may also exist. In this regard, our finding that PDE9 localizes to mitochondria whereas PDE5 is mostly in the cytosol,and that PDE9 but not PDE5 impacts mitochondrial respiration, suggests PDE9 may control this organelle by targeted post-translational changes as well. Co-localization of PDE9 to both SR membrane*(24)* and mitochondria puts it in a membrane domain within myocytes, consistent with its regulation of NP receptor-coupled cGMP. This differs from cytosolic regulation of cGMP by PDE5*(48)*. The *PDE9* gene does not contain known mitochondrial targeting sequences and its localization to this organelle may be part of a larger yet unknown protein complex. PKG also modifies proteins regulating cell growth, metabolism, and fibrosis, and these could potentially also play a role. For example, regulator of G-protein signaling 2 and 4*(49)*, transient receptor potential canonical channel type 6*(50)*, tuberous sclerosis complex protein 2*(51)* are all phosphorylated by PKG, suppressing Gq-protein coupled, NFAT/calcineurin, and mTORC1 signaling respectively. The latter can also shift metabolism to favor FA oxidation. These pleotropic effects are attractive for treating a syndrome such as obesity-CMS and HFpEF given that renal, pulmonary, vascular, metabolic, and cardiac disease commonly co-exists*(11)*.

The metabolic effects of PDE9-I documented here are compatible with its modulation of NP-derived cGMP*(24, 28)* and the demonstrated efficacy of NP to stimulate fat catabolism*(14, 52-55)* by enhancing FA oxidation. From a therapeutic perspective, however, PDE9-I offers some advantages over exogenous NP. First it is a small molecule as opposed to a peptide. Second, obesity blunts NP signaling in animals and humans by increasing expression of the NP-clearance receptor (NPRC, NPR3) in adipose tissue *(56)* while reducing that of primary NP signaling receptors*(57)*. However, PDE9 inhibition can still augment cGMP as long as there is some upstream stimulation. Consistent with this, prior studies required gene deletion of NPRC to achieve weight reduction from exogenous NP*(41)*, but this was not needed for PDE9-I. Lastly, PDE9-I augments cGMP only in those cells in which it is expressed, whereas NP and other cGMP stimulation methods activate the pathway broadly. To date, its inhibition has induced little to no change in systemic artery pressure in preclinical*(25, 26)* and clinical trials*(30)*.

There are other approaches being studied for countering obesity, all already in clinical use, mostly for diabetes. PPARα agonists such as fenofibrate and pemafibrate reduce body weight and notably triglycerides in DIO in mice*(58)*. Human studies have not shown weight loss, and their primary indication remains reducing high triglyceride levels. Fibrates activate PPARα protein so their impact depends on differential tissue expression. This is normally greatest in the liver, likely underlying their primary impact on triglycerides. As we show here, PDE9-I works differently by activating PPARα mRNA abundance in multiple tissues. This may explain the marked impact on reducing fat depots versus changes reported with PPARα-agonists in similar DIO models*(36)*. Several anti-diabetic therapies are currently being tested for weight loss. Glucagon-like peptide-1 increases insulin secretion, enhancing glucose uptake and storage as glycogen. Its anti-obesity effects are thought to be primarily due to appetite suppression with increased satiety*(59)*. Sodium-glucose co-transporter-2 (SGLT2) antagonists treat diabetes and also improve morbidity and mortality from heart failure*(60)*. They too are being studied for weight loss indications *(61)*, and though the mechanism remains uncertain, enhanced glucose/sodium excretion may play a role. Lastly, in late 2020, an FDA advisory committee overwhelmingly supported use of the combination of sacubitril and valsartan (neprilysin inhibitor and AT-1 receptor blocker) to treat HFpEF. If approved by the FDA, this would become the first approved therapeutic for HFpEF based on clinical-trial evidence *(62)*. A major mechanism of its benefit over that of AT-1 blockade is attributed to suppression of NP proteolysis by neprilysin*(63)*. Yet studies have not reported changes in PPARα signaling nor reduction in body weight*(64)* or fat transcriptomes*(65)*. This could be a matter of dose limitations to avoid blood pressure decline, and/or enhancing the signaling in the right tissues – something PDE9-I might provide in a synergistic manner.

Our study has several limitations. We used PDE9 pharmacological inhibition rather than gene deletion to test its role, as we wanted to generate substantial obesity and CMS before applying the intervention. Results using a global *PDE9* KO mouse subjected to DIO are being presented in a separate study and also reveal less weight gain and metabolic improvement over time*(66)*. While our model of obesity-CMS provides insights into HFpEF, it is not a model of HFpEF as the pressure-load was mild and mice did not develop heart failure. It is a reasonable model of obesity-CMS, incorporating the two-hit approach of DIO plus cardiac pressure-load sufficient to induce pathological hypertrophy. Notably, the severity of obesity generated in females and similar weight gain in OVX and males is lacking in most prior studies. We did not show direct evidence for PDE9-I increased PKG activity as this remains hard to detect despite cGMP increase from this regulatory pathway, likely due to a membrane-localized nano-compartment in which the signaling transpires that is lost with cell/tissue lysis. Still, the molecular signatures in the myocardium are very consistent with PKG activation. Lastly, the ChipSeq analysis is preliminary in that how the two transcriptional regulators interact remains unknown, as does whether this occurs similarly in adipocytes and other cells. However, we could find no prior similar analysis, and our results do show a striking interaction consistent with the sexual dimorphism of *Ppara* gene regulation we observed.

When first discovered, PDE9 expression was most readily detected in the brain, and based on this, all of the early pharmacological development of selective inhibitors was targeted to neurocognitive diseases such as schizophrenia and Alzheimer’s. While effective human translation remains elusive for these indications, preclinical data in the heart as since spawned clinical trials testing the impact on heart failure. This work continues, but the safety and tolerability of multiple PDE9 inhibitors has been confirmed and has negligible impact on arterial blood pressure or heart rate. Given this, the present results suggest translation to humans with obesity and CMS is feasible, and worth pursuing.

## Materials and Methods

Detailed material and methods are provided in Supplemental Materials. All statistical tests used for each data set are identified in each figure legend, along with the number of biological replicates for each group. Non-parametric models were used if a group within the experiment contained 6 or less replicates, or the distribution was not normally distributed. P-values for statistical testing are shown in individual panels or legends and explained for each.

## Supplemental Materials

### Materials and Methods

**Figure S1. Model of severe obesity and cardiac pressure-load (CMS) in female C57BL6/N mice**.

**Figure S2. Hepatic steatosis in OVX and male mice with ob/CMS model**.

**Figure S3. PDE9-I reduces obesity and improves metabolic profile in ob/CMS male mice**.

**Figure S4. PDE9-I increases myocardial cGMP, and improves male ob/CMS mouse cardiac function, pathological hypertrophic/fibrotic molecular signature, and fat metabolome**.

**Figure S5. Impact of PDE9-I on mitochondrial volume density and lipid accumulation**.

**Figure S6. PDE5A localizes to cytosol, whereas PDE9 localizes to mitochondria**.

**Figure S7. PDE9-I +/− PPARa-I does not alter food intake or activity. Non-OVX females display no significant changes in serum lipids, iWAT or gWAT weight, food intake, activity, or whole-body metabolism, or hepatic lipid accumulation in response to PDE9-I**.

**Supplemental Data Table. Clinical Features of two HpPEF groups defined by relative ratio of *PPARA* and *PDE9* gene expression**.

## Acknowledgements

The authors thank the Rodent Metabolism Core at the Center for Metabolism and Obesity Research (JHUSOM) for indirect calorimetry and GTT, Phenotyping Core for MRI body composition analysis, Small Animal CV Phenotyping and Models Core for echocardiography imaging and analysis and pressure-overload surgery, JHU Imaging Core for electron microscopy slide preparation, JHU Deep Sequencing and Microarray Core Facility for sequencing. The study was supported by: NIH R35-HL135827, RO1-HL-119012, P01HL10715, and AHA 16SFRN28620000 (DAK), NIH DK084171 (GWW), NIH-RO1-HL134821 (BOR), NIH T32-HL007227 (VSH), 16SFRN27870000 (KS), DK116625, AHA 16SFRN28620000 (SC), 16SFRN28420002 (DDS), F32 DK-116520 (RPC), and OE 688/1-1 (CUO). PF-04447943 was provided by Pfizer Inc. under a material transfer agreement.

