## Supplemental Methods, Figures, and Table for "PDE9 Inhibition Activates PPARα to Stimulate Mitochondrial Fat Metabolism and Reduce Cardiometabolic Syndrome"

**Supplemental Materials and Methods, Figures, and Tables**

### Materials and Methods:

#### Obesity/CMS Model

C57BL6/N male or female mice were fed standard chow or 60% high fat diet (D12492, Research Diets, New Brunswick, NJ, USA) starting at age 5-weeks. At 7-8 weeks, female mice were randomized to bilateral oophorectomy or sham surgery, and all mice then continued on their respective diet for ~4 months. A baseline echocardiogram was obtained, and then mice underwent mild transverse aortic constriction (mTAC) using a 26g needle. The goal was to subject the heart to modest pressure stress to stimulate some NP release. One week after mTAC, mice were then randomized to placebo (vehicle) or to the selective PDE9 inhibitor PF-04447943 (40 mg/kg/day delivered in their oral chow (yields 10 nM free plasma concentration(*1*)) or to vehicle for an additional 2 months. In other studies, mice receiving PDE9-I were further randomized to receive a PPARa inhibitor (GW6471, Cat. No. 4618, Tocris Biosciences, USA) dissolved in 1% DMSO, 95% Saline, 4% polyethylene glycol 400, 3mg/kg i.p. every other day) or the vehicle. The drug treatment protocol continued for 6-8 more weeks with serial echocardiography, indirect calorimetry, total body fat/lean mass, and serum glucose and lipids data obtained. At terminal study, tissues were harvested for histologic and molecular biology, flash frozen in liquid nitrogen and stored at -80°C until analyzed. Additional studies used *PDE9* global KO mice and WT littermate controls(*2*) maintained on the same HFD starting age 5 weeks and continued for 26 weeks.

**Conscious Mouse Echocardiography:** was performed in conscious mice using a Vevo 2100 (VisualSonics) with an 18-38 MHz transducer (SanoSite Inc.), and images analyzed with VisualSonics image software. The sonographer was blinded to genotype and treatment. Ejection fraction and estimated wall mass were derived using standard methods (*2*).

#### Blood collection and analysis

Blood samples were collected by tail vein bleeding, and serum isolated using Microvette CB 300 (Sarstedt, Newton, NC, USA). Serum triglyceride and cholesterol were measured according to the manufacturer's instructions using Infinity kit (Thermo Fisher Scientific, Middletown, VA, USA).

**Glucose tolerance test**

Mice were subjected to overnight food deprivation (approximately 16 hours), after which glucose (1 mg/kg) was injected intraperitoneally and baseline blood glucose measurement was taken from a clean tail blood droplet using a glucometer (BD Logic, NovaMax strips, Cat 8548043523). Repeat measurements were made at 15, 30, 60, and 120 min.

**Magnetic Resonance Imaging for body composition analysis**

Total body fat and lean mass composition was determined by Echo-Magnetic Resonance Imaging (MRI)-100 (Echo Medical Systems, Waco, TX, USA) by the JHSOM Mouse-phenotyping Core using standard protocols.

**Indirect calorimetry, caloric intake, and physical activity measurements**

Indirect calorimetry and additional measures of food intake and physical activity were performed in open-circuit indirect calorimetry cages (Comprehensive Lab Animal Monitoring System, Columbus Instruments, Columbus, OH,USA). Mice were monitored individually and continuously for 4 days: 2-3 days to monitor adaptation to the novel environment, and the third or fourth day for reporting data. The system periodically measured rates of $O_2$ consumption ($VO_2$) and $CO_2$ production ($VCO_2$) for each cage, as well as food intake and physical activity (infrared beam array). Data were processed in time segments (daily 24-hr, and 12-hr dark and light), and averages calculated for each mouse in each segment. $VO_2$, $VCO_2$ were normalized to 0.8 x lean mass + 0.2 x fat mass as described(*3*), using data from EchoMRI-100 acquired within a few days of indirect calorimetry.

**Isolated myocyte Studies:** Rat neonatal cardiomyocytes were isolated from newborn pups and cultured for 2 days prior to study.

***Study 1: Lipid loading and impact of PDE9-I*:** Cardiomyocytes were incubated with 2.5 µM PF-7943 or vehicle for 24 hrs prior to exposure to FA (mixture of oleic (Cat O1008, Sigma-Aldrich, USA), linoleic (Cat L1376, Sigma-Aldrich, USA) and palmitate acid (Cat P0500, Sigma-Aldrich, USA) complexed (3:1 ratio) to a 3% fatty acid free bovine serum albumin (BSA) at a concentration of 0.4 mmol/L. After an additional 24 hours incubation, cells were fixed in

4% formaldehyde for 30 min, washed with PBS x3 and stained with Oil Red O. After gentle washing with PBS x3, cells were imaged using phase-contrast microscopy (Zeiss AxioObserver A1 inverted microscope & Olympus DP80 dual RGB and monochrome camera) to identify lipid-laden vesicles. Images were processed with ImageJ to determine vesicles/cell. Alternatively, the cells were washed with PBS, loaded with 5 μM BODIPY 493/503 (D3922, Invitrogen, USA) for 20 min at 37 °C, fixed in 4% paraformaldehyde, washed, permeabilized and counterstained with 2-(4-amidinophenyl)-6-indolecarbamidine dihydrochloride (DAPI) (D1306, Invitrogen, USA). Images were taken using Revolve microscope from Echo Laboratories (San Diego, CA, USA) and analyzed using ImageJ (v1.52) software after background subtraction and threshold detection.

***Study 2: Co-localization of GFP-labeled PDE9 and mitotracker Red.*** NRCMs were infected with adenovirus (AdV) containing a *Gfp-PDE9* (MOI 10), and after 48 hours, cells stained with 200 nmol/L Mitotracker Red CMXRos (M7512, Invitrogen, USA) to label mitochondria, and imaged with confocal microscopy (Zeiss AxioObserver A1 inverted microscope & Olympus DP80 dual RGB and monochrome camera).

***Study 3: Effect of PDE9 versus PDE5A on mitochondrial oxygen consumption and ATP generation.*** Mitochondrial respiration was determined from oxygen consumption rate (OCR) measured by a Seahorse XF96 Extracellular Flux Analyzer (Seahorse Biosciences, MA, USA), per manufacturer's protocol. NRCMs were isolated from 2–3-day-old neonatal Sprague-Dawley rats. 40,000 cells were seeded into fibronectin-coated wells of a Agilent Seahorse XF96 well cell culture plate in 10% FBS containing DMEM media. 24 hrs later, a confluent monolayer of spontaneously beating NRCMs was formed. For one study, NRCMs were treated with or without PDE9-I (2 and 5 μM) overnight, media then replaced with DMEM+Fatty acid mixture as noted above + 0.5 mM carnitine and cultured for an additional 24hrs. In the other study, cells were infected with Adenovirus expressing *Gfp, PDE9-Gfp* (C-terminus, MOI 10), or *Pde5a-Gfp* (C-terminus, MOI 10) and incubated for 48 hrs followed by OCR measurements. To assess OCR, cells were tested in DMEM without phenol red supplemented with sodium pyruvate (pH-7.4). After calibration of the sensor cartridge, the XF96 plate was placed into the seahorse instrument and OCR was measured as pmole per minute per cell. Analysis of mitochondrial function was made using changes in OCR in the presence of 1.0 μM oligomycin (inhibits complex V – basal

OCR determined), 1.0 µM FCCP (uncoupling agent revealing maximal OCR through complex IV), 0.5 µM Rotenone and Antimycin A (inhibits complex 1 shutting down mitochondrial respiration to calculate non-mitochondrial respiration). After each injection, the seahorse instrument measured OCR for four times at five time points. The results were normalized for cell protein in each well and averaged.

**Lipolysis Assays**

*Lipolysis in differentiated adipocytes:* 3T3-L1 preadipocytes from Zen-Bio, Inc (KT-01, USA) were grown and differentiated per manufacturer's instructions. Differentiated adipocytes were maintained in 3T3-L1 adipocyte maintenance medium and incubated with vehicle (0.1%DMSO), PDE9-I (2.5 µM PF-7943), or 3-isobutyl-1-methylxanthine (IBMX, 100 µM) or isoproterenol (1 µM) as positive controls for 24hrs. Induction of lipolysis was assessed by glycerol release using adipocyte lipolysis assay kit (LIP-1-NC, Zen-Bio, Inc) per manufacturer's protocol.

*Lipolysis analysis in Hep G2 and Neonatal Cardiomyocytes:* Hep G2 cells (Human, ATCC cell line HB-8065) were seeded at a density of $5\times 10^3$ cells/well and NRVMs were plated at a density of 40000/well into 96-well plates. Cells were maintained in a medium containing the FA mixture (see above) and treated with combinations of the following compounds for 24 hrs: ANP (10 µM, Cat S-20647, Anaspec), PDE9-I (2.5 µM or 5µM PF-7943), and PPARa inhibitor GW6471 (10 µM, Cat 4618, Tocris). In each case, cells were first incubated with the respective inhibitors for 3hrs prior to ANP stimulation. Another group of NRVMs were transfected with either scrambled control siRNA (25 nM, Cat D-001810-10) or PDE9 Rat targeted siRNA (25 nM, Cat L-098890-02) from Horizon Discovery.

**PPARα promoter activity assay**

Hep G2 (Human, ATCC cell line HB-8065) cells were cultured in DMEM (10% FBS, 1% penicillin/streptomycin, Thermo Fisher) to 70% confluence. Cells were transfected using Xfect transfection reagent ( Cat. No. 631317, Takara Bio, USA, ) with recombinant plasmid containing 1206 bp upstream TSS of human PPARa promoter-luciferase reporter (0.3 µg) (generously provided by Dr. Bert Staels, University of Lille, France). Renilla-luciferase plasmid (0.001 µg; Promega) was transfected as an internal control. After 24 hrs, cells were incubated with 8-pCPTcGMP (C5438, Sigma-Aldrich, USA) at varying concentrations for 6 hours. Cell lysates were then extracted using passive lysis buffer (E1910, Promega Corp., USA), and luciferase activity measured by Dual-Luciferase Reporter Assay (E1910, Promega Corp., USA) using a GloMax96 plate reader (Bio Tek, USA).

**Mitochondrial isolation and Protein analysis**

NRCMs were infected with AdV expressing *Pde5a-Gfp* or *PDE9-Flag* (each at 10 MOI) and maintained in DMEM with 10% FBS. After 48 hours, cells were scraped into ice cold PBS and spun at 800 x g for two minutes, and mitochondria isolated using a kit per manufacturer's instructions (Thermo Scientific, Cat 89874, Lot TJ272027). Mitochondria were reconstituted in SDS sample loading buffer (Licor, Cat 928-40004), run on 4-20% Mini-Protean TGX gels (BioRad), and blotted onto a nitrocellulose membrane (BioRad). The following antibodies were used: alpha-Tubulin (Cell Signaling, 3873S, Lot 12), GFP (Invitrogen, A6455, Lot 2185052), Flag (Sigma, F1804, Lot SLBW5142), VDAC1/Porin (Abcam, ab15895, Lot GR3237200-1), and Total Protein Stain (Licor, 926-11016). Antibody binding was visualized with an infrared imaging system (Odyssey, Licor) and quantification by Odyssey Application Software 3.1.

**Light microscopy histology**

Immediately after removal, liver, white (inguinal and gonadal), and brown adipose tissues were fixed overnight by immersion at 4°C in 4% paraformaldehyde. Paraffin-embedded dewaxed 4 μm sections were stained using hematoxylin and eosin.

**Transmission Electron Microscopy and Immunogold Staining**

Freshly extracted Heart tissues were fixed in 2.5% glutaraldehyde, 3mM $MgCl_2$, in 0.1 M sodium phosphate (Sorenson's) buffer, pH 7.2 for one hour at room temperature. After buffer rinse, samples were postfixed in 1% osmium tetroxide, 0.8% potassium ferrocyanide in 0.1 M Sorenson's buffer (1 hr) on ice in the dark. Following a 0.1 M maleate buffer rinse, samples were stained with 2% uranyl acetate (0.22 μm filtered, 1 hr in the dark) in maleate buffer, dehydrated in a graded series of ethanol, followed by propylene oxide and embedded in Eponate 12 (Ted Pella, Inc.) resin. Samples were polymerized at 60 °C overnight. Thin sections, 60 to 90 nm, were cut with a diamond knife on the Reichert-Jung Ultracut E ultramicrotome and picked up with 2x1 mm formvar coated copper slot grids. Grids were stained with 2% uranyl acetate in 50% methanol and 0.4% lead citrate before imaging on a Hitachi 7600 TEM at 80 kV. Images were captured with an AMT XR50 CCD (5 megapixel) camera at Johns Hopkins Microscopy Core. Mitochondrial density and area measurements were performed using ImageJ (v1.52). Scales were set based on the 500 nm legend for each image. The adjust threshold function was used to highlight mitochondria and the measure function to determine pixel intensity and area of the highlighted region.

For immunogold staining, NRVMs were plated in a 6 well plate (1X $10^5$ cell per well). 24 hours later, cells were transfected with either control GFP adenovirus (AdV) or GFP-tagged-*PDE9* adenovirus (MOI 10). 48 hrs after transfection, cells were fixed in EM grade 4% paraformaldehyde, 0.1% glutaraldehyde in 80 mM phosphate buffer (Sorenson's buffer) with 3 mM magnesium chloride, pH 7.2. All subsequent steps were done at $4^0$C until 70% ethanol dehydration. Samples were rinsed 3 times in buffer containing 3% sucrose, then incubated in 1.5% potassium ferrocyanide reduced 1% osmium tetroxide in 80 mM phosphate buffer, containing 3 mM magnesium chloride followed by two distilled water rinses, dehydrated in a graded series of Ethanol, followed by propylene oxide, and embedded in EPON (Ted Pella, Inc.). Plates are baked at $37^0$C for 2-3 days before an overnight bake at $60^0$C. 80-90 nm ultra-thin sections were picked up on formvar coated 200 mesh nickel grids and allowed to dry. Grids were floated on all subsequent steps. All solutions were filtered except for antibodies which were centrifuged at 13K for 5 min. Grids were wetted 3 times on D-$H_2O$, then placed on 3% sodium metaperiodate for 20 min. After a 15 min rinse in D-$H_2O$, grids were placed in 50 mM $NH_4Cl$ in TBS for 10 min, followed by 20 min block in blocking solution (3% normal goat serum and 3% bovine serum albumin in TBST). Antibody (anti GFP, Abcam ab6556) incubation was done at appropriate dilutions (1:100) in the same block. Antibody block (no primary) served as negative control. Incubations were carried out at $4^0$C overnight. Blocks were equilibrated to room temperature, grids were placed on blocking solution for 10 min, followed by a 1 min rinse in TBS. Gold conjugated secondary antibody (12nm, Colloidal gold 109-185-088, Jackson Immuno Research Labs) was diluted 1:40 in TBS and sections were incubated for 2 hrs at room temperature in a humidity chamber. After a 10 min TBS incubation followed by a quick D-$H_2O$ rinse, grids were hard fixed in 1% glutaraldehyde in 100 mM sodium cacodylate buffer for 5 min. After a brief D-$H_2O$ rinse, grids were stained with 2% uranyl acetate (aq.) for 20 min, rinsed again with D-H20; blot dried and allowed to dry overnight before viewing. Images were captured at 120 kV on a Ceta Camera (8 Mpixel CCD, 16bit 40fps) equipped on Talos L120C G2 Transmission Electron Microscope at Johns Hopkins SOM Microscopy Core.

**RNA isolation and Gene expression analysis**

Total RNA from Heart, WAT, and BAT were extracted using Trizol Reagent (Cat. No. 15596026, Invitrogen, Thermofisher, USA) per manufacturer's instructions. High-Capacity RNA-to-cDNA Kit (Cat. No. 4388950, Applied Biosystems, Thermofisher, USA) was used to reverse transcribe the RNA into cDNA. The PCR-array gene expression profiling was performed using the Bio-Rad Prime PCR plates dedicated to Carbohydrate (10029626, M384) and Lipid metabolism (10040365, M384) gene panels according to manufacturer's instruction.

Quantitative real time PCR analysis was carried out using TaqMan specific primers: Acadm (mouse # Mm01323361_mH), CD36(mouse # Mm00432403_m1), Cpt1a (mouse # Mm00550448_m1), Hadh (mouse # Mm00492535_m1), Acot1 (mouse # Mm01622471_s1), PDK4 (mouse # Mm01166879_m1), Dgat1 (mouse # Mm00515643_m1), fabp1 (mouse # Mm00444340_m1), Ucp1 (mouse # Mm01244861_m1), Cidea (mouse # Mm00432554_m1), Prdm16 (mouse # Mm00712556_m1), Dio2 (mouse # Mm00515664_m1), Adrb3 (mouse # Mm02601819_g1), PPARα (mouse # Mm00440939_m1), Atp6 (mouse # Mm03649417_g1), cox1 (mouse #Mm04225243_m1), cycs (mouse #Mm01621048_s1), Nd1 (mouse # Mm04225274_s1), Nrf1 (mouse # Mm00447998_m1), ppargc1a (mouse # Mm01208835_m1), Tfam1 (mouse # Mm00447485_m1), Nppa (mouse # Mm01255747_g1), Ctgf (mouse # Mm01192933_g1), Postn (mouse # Mm01284913_g1), Tgfb (mouse # Mm01178820_m1), Col1a1 (mouse # Mm00801666_g1), Lox (mouse # Mm00495386_m1) or gylceraldehyde-3-phosphate dehydrogenase (GAPDH, mouse #99999915_g1,) by Applied Biosystems. The threshold cycle (Ct) values were determined by crossing point method and normalized to GAPDH (Applied Biosystems) values for each run.

**Cyclic GMP assay**

Tissue cGMP levels were determined using an EIA assay (Cyclex, USA) as described (*2*).

**Metabolic profiling of BAT and Myocardial Tissue for Acylcarnitines**

Upon sacrifice, five replicates of each experimental series of heart and BAT tissues were washed for 5s in ice-cold PBS to remove excess blood and then snap-frozen in liquid nitrogen. Tissues were ground in liquid nitrogen. 15 mg of frozen ground tissue powder was further homogenized using ice -cold extraction solution (40% acetonitrile/40% methanol/20% water, spiked with internal standards). Homogenized samples were then incubated at -20 ˚C for 1 h. After incubation, all the samples were centrifuged at 16K for 15 min at 4°C. Supernatants were transferred to new glass tubes and dried using speed vac. Pellets were used for protein estimation using the BCA method.

Extracted myocardial tissue and brown adipose tissue samples were assayed using stable-isotope dilution ($^{2}H_{3}$-carnitine (d3C0), $^{2}H_{3}$-acetyl- (d3C2), $^{2}H_{3}$-propionyl- (d3C3), $^{2}H_{3}$-butyryl- (d3C4), $^{2}H_{9}$-isovaleryl- (d9C5), $^{2}H_{3}$-octanoyl- (d3C8), $^{2}H_{9}$-lauroyl- (d9C12), $^{2}H_{9}$-miristoyl- (d9C14), $^{2}H_{3}$-palmitoyl- (d3C16) and $^{2}H_{3}$-stearoylcarnitine (d3C18). Cambridge Isotope Laboratories (CIL) NSK-B and NSK-B-G1) after were reconstituted in 0.5mL 10% methanol in water + 0.1% formic acid by means of 10 minutes of orbital shaking and thorough vortexing.

112 species comprising aliphatic acylcarnitines ranging from C0 (free) to docosaenoyl- (C22), and including odd-chain, branched-chain, unsaturated, hydroxylated and dicarboxylic moieties, were analyzed using a Waters Acquity UHPLC in a ACE C18-PFP column (2.1 x 100 mm, 3µm particle-size) at 0.3mL/min at 30°C with water partitioning in 3:1 acetonitrile:methanol (by volume), both + 0.1% formic in gradient elution (5 min, 0%B; 8 minutes to 23%B and then to 100%B in 18 minutes) and monitored in a Sciex 4500 triple quadrupole mass spectrometer using optimized specific transitions to class specific product ion m/z 85 at defined retention times. They also had monitored the neutral loss of trimethylamine from the parent ion fragment, as a confirming transition. Injection volume was 5µL.

Species were quantified using 6 non-zero levels calibration curves using the authentic standard (CIL, NSK-B-US and NSK-B-US-G1, where possible, spiked in analyte stripped serum (Biocell Laboratories); if not available, the closest in the homologue series. Unsaturated species were assayed using the same response factor than the corresponding aliphatic species of the same chain length. Hydroxylated were referred to hydroxyisovaleryl- (C5OH) or hydroxylpalmitoylcarnitine (C16OH) and dicarboxylic to glutarylcarnitine (C5DC). Replicates showed acceptable coefficients of variation (20%).

**Human HFpEF and control myocardial biopsy analysis**

As previously reported(*4*), RNAseq was performed on 41 flash frozen 2-3.5 mg right ventricular septal endo-myocardial biopsies obtained from HFpEF patients referred to the Johns Hopkins University HFpEF Clinic from 1/2016-4/2018. Each patient provided informed consent for the research protocol which was approved by the Johns Hopkins Institutional Review Board (IRB). Patients were phenotyped by clinical assessment, echocardiography, and invasive hemodynamics, with HFpEF diagnosis based on signs and symptoms of clinical HF, LV ejection fraction ≥ 50% by echocardiography within prior 12 months, and at least two of: 1) increased left ventricular [LV] wall thickness, left atrial [LA] diameter) or diastolic dysfunction on echocardiography; 2) N-terminal pro–B-type natriuretic peptide (NTproBNP) ≥ 125 pg/mL; or 3) pulmonary artery wedge pressure ≥ 15 mmHg at baseline; or ≥ 25 mmHg with exercise(*4*). Patients with EF<50%, greater than moderate valvular disease, infiltrative cardiomyopathy (including cardiac amyloidosis), restrictive cardiomyopathy, congenital heart disease, constrictive pericarditis, isolated pulmonary arterial hypertension, hypertrophic cardiomyopathy (known genetic variant, severe unexplained LVH, or presence of myocyte disarray on histology), or prior heart transplantation were excluded. RV mid-septal myocardium from explanted but unused donor control hearts (CON, n=24) provided by the University of Pennsylvania under an IRB-approved protocol served as the comparison group. The human genome was obtained in FASTA format (GRCh38) from Ensembl version 92 and gene set annotation in gtf format. The hisat2 indices were built from the genome index using hisat2-build (*5*) from Hisat2 version 2.1.0. Raw RNAseq paired-end reads were aligned to the genome using hisat2 (default flags). The total reads per sample ranged from 30-50 million and all sample alignment mapping rates were above 97%. HTSeq was used to count reads mapping to individual genes by processing the sorted bam files with accepted read quality. (*6*) Differentially expressed genes were defined using the 5% FDR (Benjamini-Hochberg) threshold for significance. Gene pathway enrichment (KEGG) and gene ontology (GO) was determined using clusterProfiler using R.

**Chromatin Immunoprecipitation Sequencing (ChIP Seq) Analysis of PPARα binding and influence of ERα and ERβ co-activation**

ChIP was performed in Hep G2 cells with two independent full biological replicates. Approximately $5 \times 10^6$ Hep G2 cells (Human, ATCC cell line, HB-8065) were cultured into three 15 cm diameter plates in DMEM with 10% fetal bovine serum. After 24 hours, cells were transfected with expression vectors for either PPARa alone (Human Tagged ORF, RC216176, Origene), or combined with estrogen receptor alpha (ERa) (pCMV ERalpha, #101141, Addgene) or estrogen receptor beta (ERb) (pcDNA Flag ERbeta, #35562 Addgene). 48 hrs after transfection, cells were either treated with a PPARa agonist alone for 2hrs (10 µM, Wy14643, Cat 1312, Sigma-Aldrich) or first with an ERa agonist (10 nM, PPT, Cat 1426/50, Tocris Biosciences, USA) or ERb agonist (10 nM, DPN, Cat 1494/50, Tocris Biosciences, USA) for 45 mins, followed by 2 hrs incubation with the PPARa agonist. SimpleChIP Enzymatic Chromatin IP kit (Cat. No. 9003, Cell Signaling Technology, US) was used for the immunoprecipitation per manufacturer's instructions. Cells were washed x2 with ice cold PBS and then crosslinked by incubating with 1% formaldehyde for 10 min. The reaction was stopped by the addition of 2.5 M glycine and the plates washed twice with ice cold PBS. Chromatin was treated with nuclease and subjected to sonication. Immunoprecipitation was performed with PPARα specific antibody (ab227074 ChIP-grade, Abcam ) and Protein G magnetic beads. Chromatin was eluted from the beads, uncrosslinked and purified. ChIP and input DNA libraries were prepared using Clontech DNA SMART kit (Cat. No. 634866, Takara Bio,USA ). ChIP-seq was performed according to the ENCODE (The Encyclopedia of DNA Elements) consortium's standards and guidelines. Sequencing was carried out by the JHU Genomics Core Facility using an Illumina NextSeq 500 sequencing system. Irreproducible Discovery Rate (IDR) was used to evaluates reproducibility of the experiments by measuring consistency between our biological replicates within an experiment. The data is provided in the GEO (submission # GSE156956). Data quality was validated using FastQC (v0.11.9). Sequenced reads were trimmed for adaptor sequence, and for low-quality sequence, using Trim Galore v0.6.4_dev. Gene Alignment was done to hg38 whole genome using bowtie (v1.0.1) and peak locations were determined using the MACS2 algorithm v2.2.7.1 (0.01 FDR cut off, fold enrichment 2, 3 and 5). Peaks were annotated using Homer v4.11 and functional analysis was performed using R (version 3.3.6), ChIPSeeker (v1.18.0, tss region -2500, 2500) and DiffBind (v2.16) packages.

**Supplemental Figures and Legends**

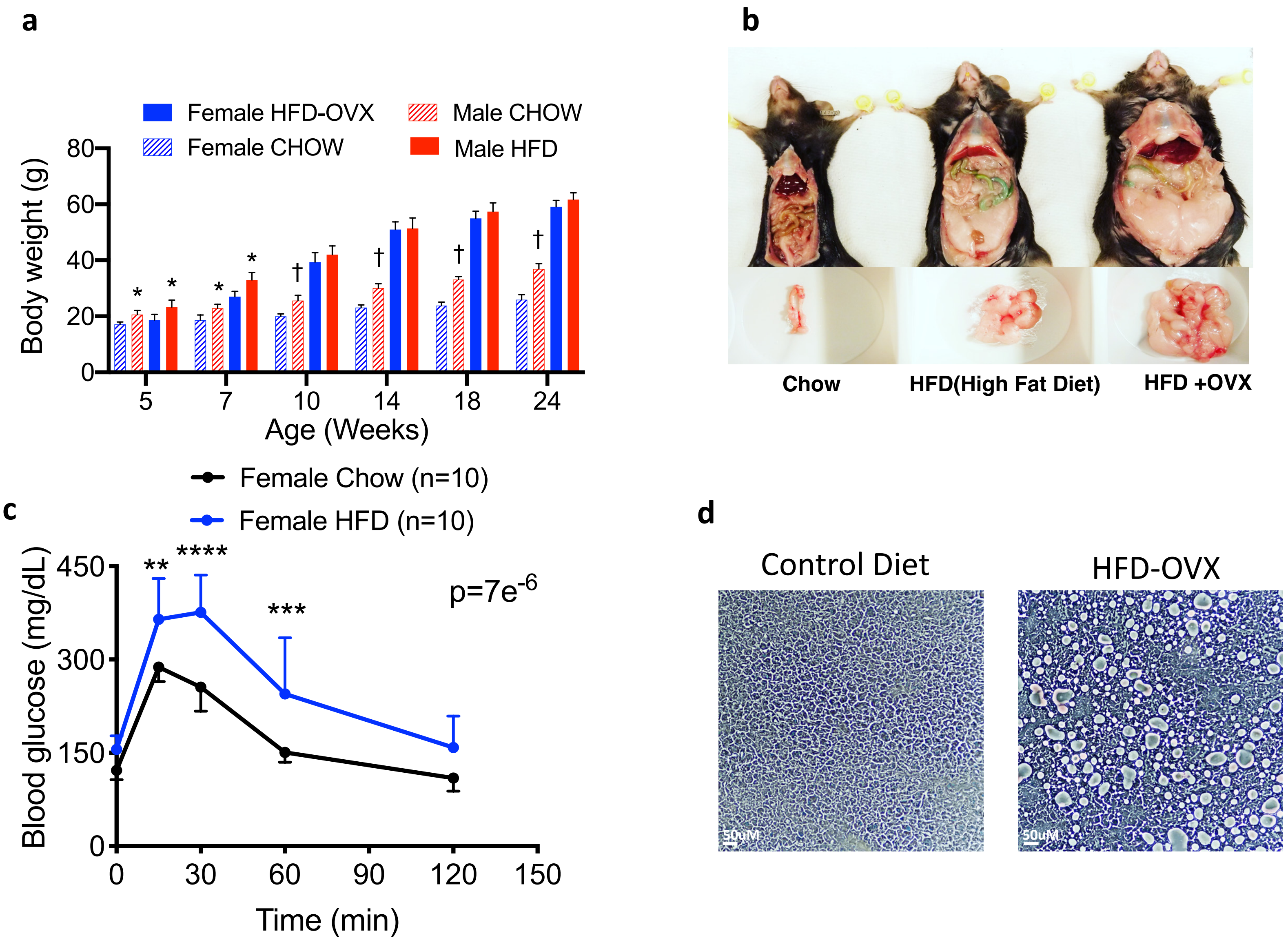
a
Female HFD-OVX
Female CHOW
Male CHOW
Male HFD
Body weight (g)
80
60
40
20
0
5
7
10
14
18
24
Age (Weeks)
b
Chow
HFD(High Fat Diet)
HFD +OVX
c
Female Chow (n=10)
Female HFD (n=10)
Blood glucose (mg/dL)
450
300
150
0
**
****
***
p=7e-6
0
30
60
90
120
150
Time (min)
d
Control Diet
HFD-OVX

**Figure S1**

**Figure S1. Model of severe obesity and cardiac pressure-load (CMS) in female C57BL6/N mice.** **a)** Total body mass increase over 6-month period for ovariectomized (OVX) females and males with control or high-fat diet. Male weights exceed females until several weeks after OVX when they become equal. At 6-months, weights in OVX females are >2x that of chow diet female controls. * p<0.005, † p<0.0005 vs same diet in females. 2WANOVA, SMCT. **b)** Example of marked abdominal adiposity in females with OVX+HFD. **c)** Depressed glucose tolerance in OVX+HFD versus female chow-diet controls (n=10/group, 2WANOVA, SMCT; ** p=0.002, *** =0.0001, and ****=6.7e$^{-7}$ between groups). **d)** Representative micrographics of liver in normal versus HFD showing marked steatosis in HFD+OVX mice measured at 13 weeks prior to when subsequent mTAC and drug treatment randomization occurs (replicated x8).

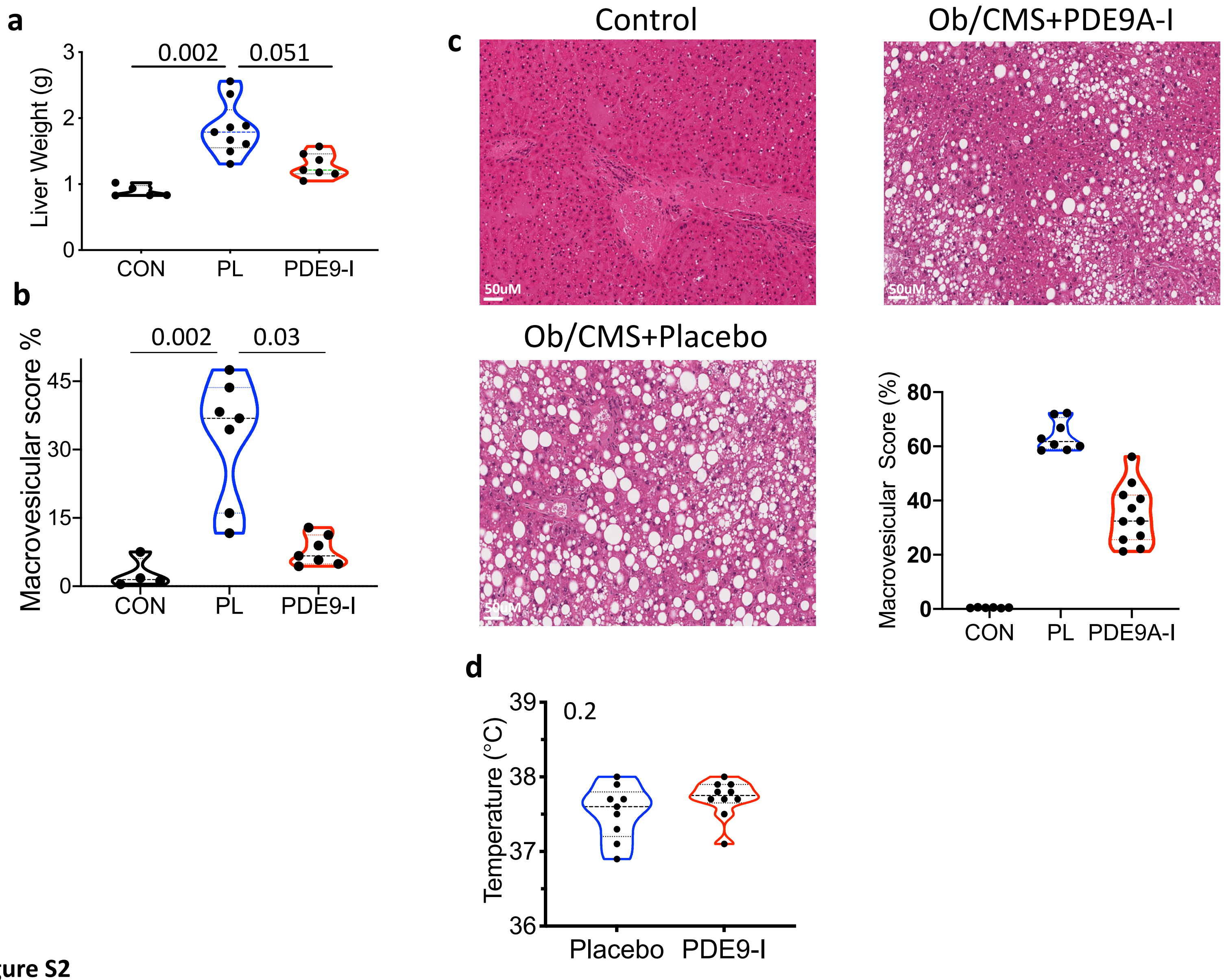

a
Liver Weight (g)
0.002
0.051
CON
PL
PDE9-I
b
Macrovesicular score %
0.002
0.03
CON
PL
PDE9-I
c
Control
Ob/CMS+PDE9A-I
Ob/CMS+Placebo
50uM
Macrovesicular Score (%)
CON
PL
PDE9A-I
d
Temperature (°C)
0.2
Placebo
PDE9-I


Figure S2

**Figure S2. Hepatic steatosis in OVX and male mice with ob/CMS model. a)** Quantification of hepatic mass and **b)** percent macrovesicular lipid accumulation in OVX mice with ob/CMS model. Related to examples displayed in Figure 1f. PL-placebo, CON- chow diet, KW with DMCT p-values shown. **c)** Example liver histology and summary analysis of macrovesicular lipid accumulation in male mice with ob/CMS model. KW with DMCT. **d)** Core temperature in OVX ob/CMS mice treated with placebo versus PDE9-I.

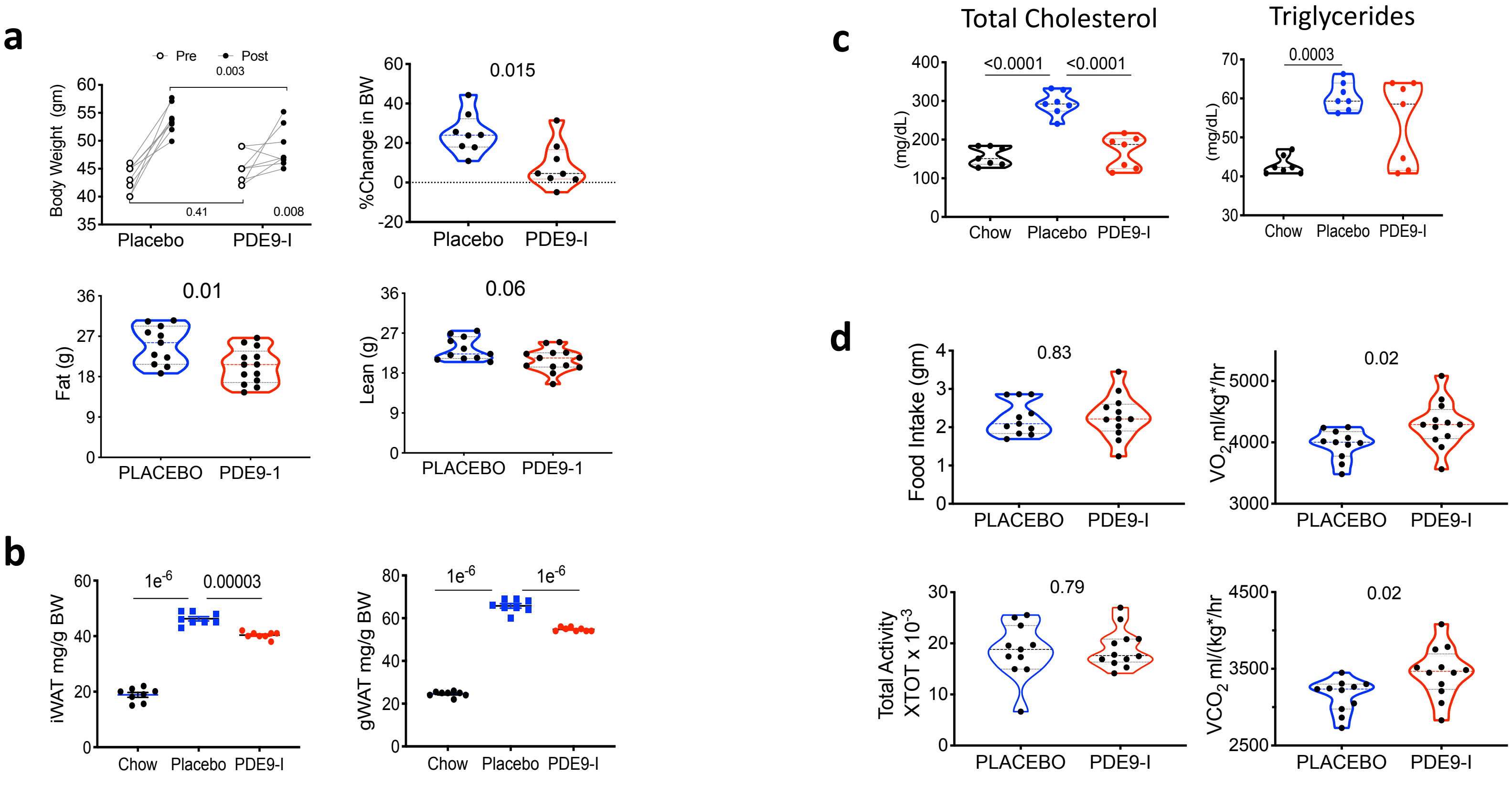
a
Pre
Post
0.003
Body Weight (gm)
60
55
50
45
40
35
0.41
0.008
Placebo
PDE9-I
%Change in BW
0.015
60
40
20
0
-20
Placebo
PDE9-I
Fat (g)
0.01
36
27
18
9
0
PLACEBO
PDE9-1
Lean (g)
0.06
36
27
18
9
0
PLACEBO
PDE9-1
b
iWAT mg/g BW
1e-6
0.00003
60
40
20
0
Chow
Placebo
PDE9-I
gWAT mg/g BW
1e-6
1e-6
80
60
40
20
0
Chow
Placebo
PDE9-I
c
Total Cholesterol
(mg/dL)
<0.0001
<0.0001
400
300
200
100
0
Chow
Placebo
PDE9-I
Triglycerides
(mg/dL)
0.0003
70
60
50
40
30
Chow
Placebo
PDE9-I
d
Food Intake (gm)
0.83
4
3
2
1
0
PLACEBO
PDE9-I
VO2 ml/kg*/hr
0.02
5000
4000
3000
PLACEBO
PDE9-I
Total Activity XTOT x 10-3
0.79
30
20
10
0
PLACEBO
PDE9-I
VCO2 ml/(kg*/hr
0.02
4500
3500
2500
PLACEBO
PDE9-I

**Figure S3**

**Figure S3. PDE9-I reduces obesity and improves metabolic profile in ob/CMS male mice. a)** Total body weight before and after treatment in placebo vs PDE9-I treatment groups. (n=8, 2WANOVA, SMCT); percent change from same data, MW; MR-derived total body fat and lean mass for both groups **(**n=11/13, MW**)** **b)** iWAT and gWAT from males in both treatment arms. (n=8/group, 1WANOVA, TMCT). **c)** End-of study serum cholesterol and triglycerides (n=7/group, 1WANOVA, TMCT). **d)** Indirect calorimetry results from same experiment (n=11/12 per group, unpaired t-test). kg* = (lean mass + fat mass * 0.2).

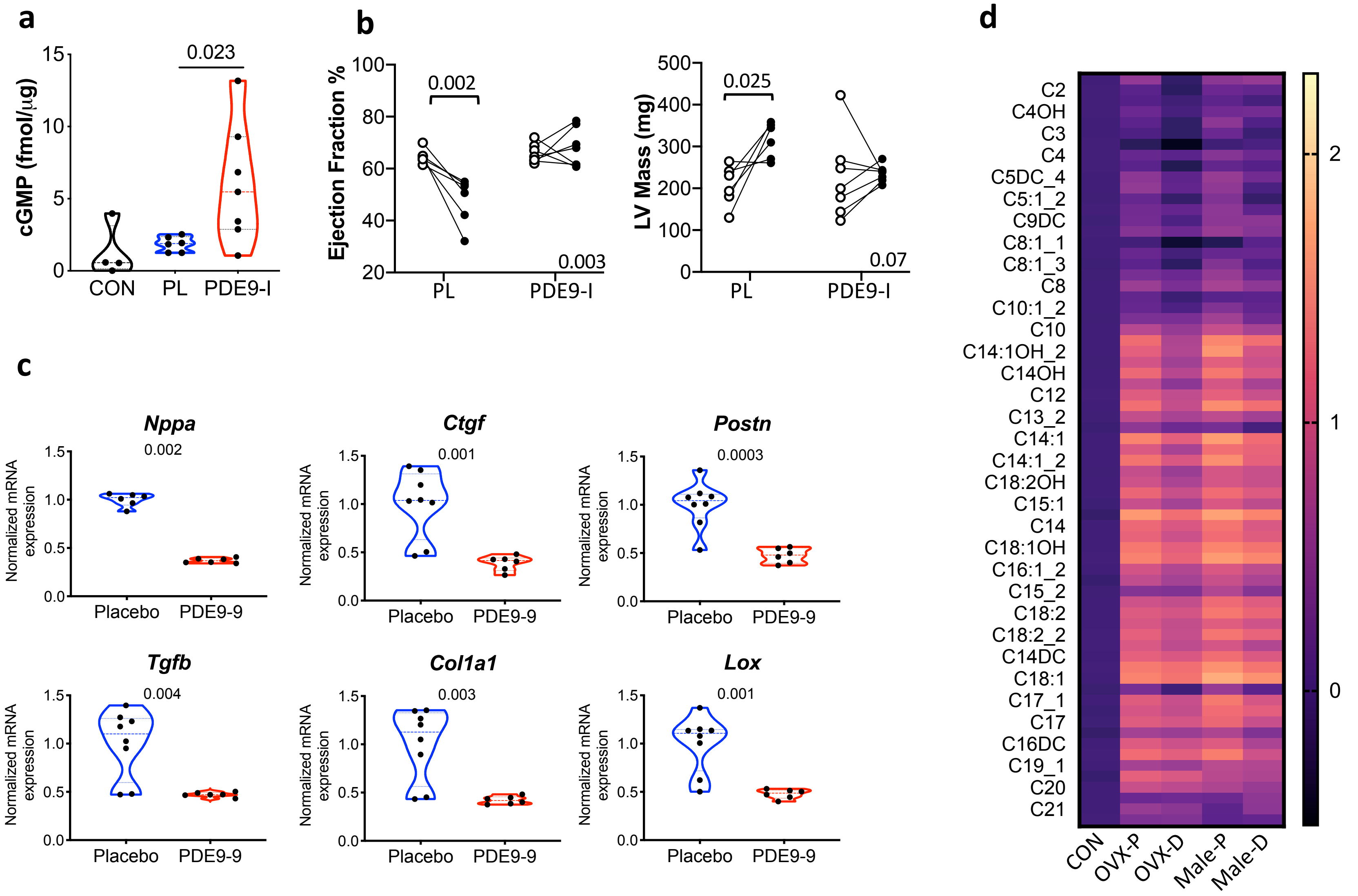

a
cGMP (fmol/µg)
0.023
CON
PL
PDE9-I
b
Ejection Fraction %
0.002
0.003
PL
PDE9-I
LV Mass (mg)
0.025
0.07
PL
PDE9-I
c
Nppa
0.002
Ctgf
0.001
Postn
0.0003
Tgfb
0.004
Col1a1
0.003
Lox
0.001
Normalized mRNA expression
Placebo
PDE9-9
d
C2
C4OH
C3
C4
C5DC_4
C5:1_2
C9DC
C8:1_1
C8:1_3
C8
C10:1_2
C10
C14:1OH_2
C14OH
C12
C13_2
C14:1
C14:1_2
C18:2OH
C15:1
C14
C18:1OH
C16:1_2
C15_2
C18:2
C18:2_2
C14DC
C18:1
C17_1
C17
C16DC
C19_1
C20
C21
CON
OVX-P
OVX-D
Male-P
Male-D


**Figure S4**

**Figure S4. PDE9-I increases myocardial cGMP, and improves male ob/CMS mouse cardiac function, pathological hypertrophic/fibrotic molecular signature, and fat metabolome. a)** Myocardial cGMP assessed in OVX mice on chow (CON) or with ob/CMS model treated with placebo (PL) or PDE9-I. KW, DMCT. **b)** Echocardiography for ejection fraction and LV mass in male ob/CMS mice treated with placebo or PDE9-I. (n=6,7/group, 2WANOVA p-value for interaction at lower right, TMCD p-values displayed). **c)** Quantitative PCR analysis of mRNA for hypertrophic and pro-fibrotic signaling genes in myocardium of males at termination of the same protocol. Abbreviations as provided in Figure 2C legend. KW test p-values displayed. **d)** Heat map (log transformed) for acyl-carnitine metabolites in myocardium of mice fed normal diet (CON) or OVX or Males with ob/CMS treated with placebo (PL) or PDE9-I. Levels increased in both groups with obesity in placebo and were significantly reduced by treatment.

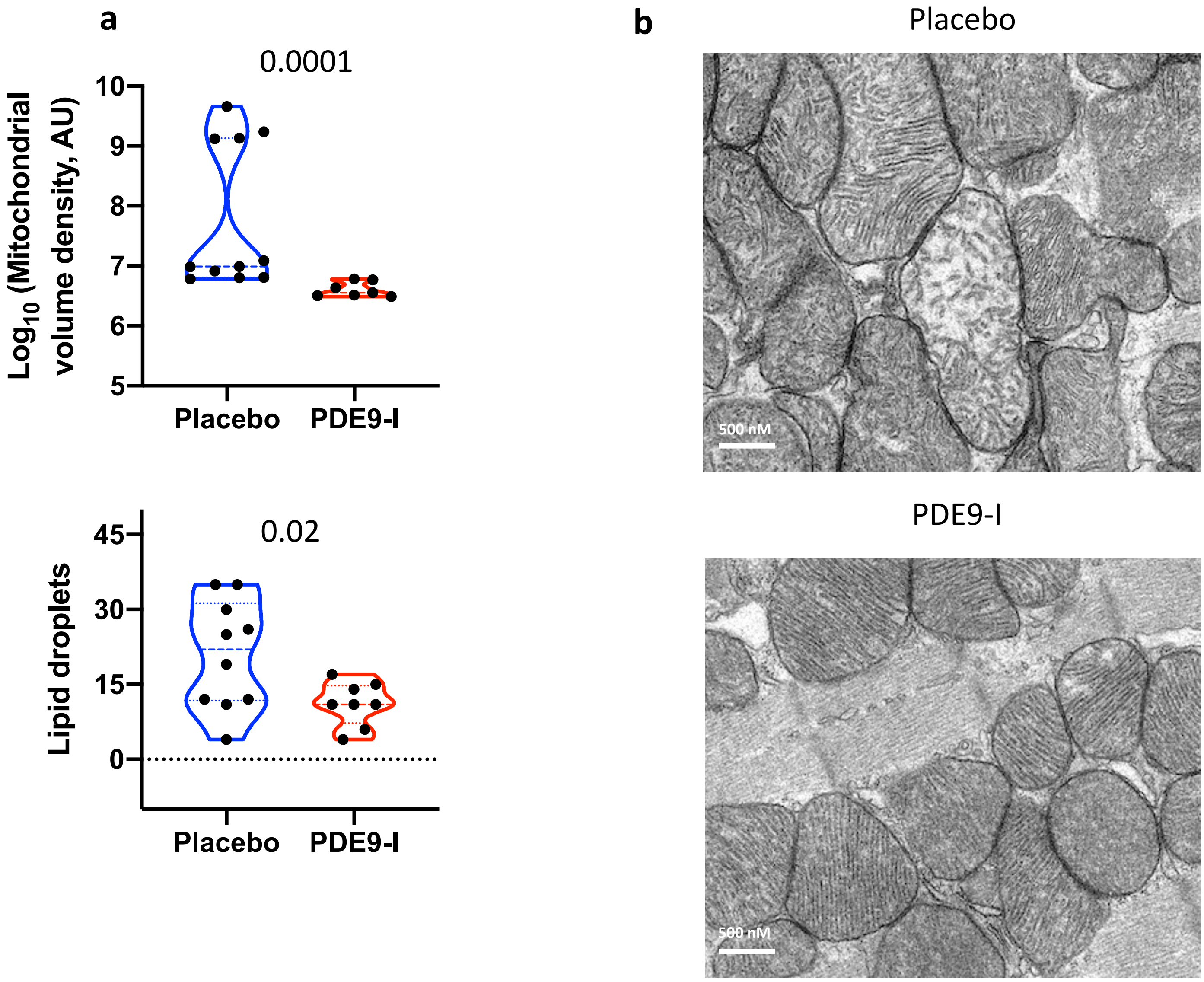

a
0.0001
Log10 (Mitochondrial volume density, AU)
10
9
8
7
6
5
Placebo
PDE9-I
0.02
Lipid droplets
45
30
15
0
Placebo
PDE9-I
b
Placebo
500 nM
PDE9-I
500 nM


**Figure S5**

**Figure S5. Impact of PDE9-I on mitochondrial volume density and myocardial lipid accumulation. a)** Quantitation of mitochondrial volume density (upper) and lipid droplets in myocardium of OVX mice with ob/CMS after placebo versus PDE9-I treatment. Relates to example images shown in Figure 4f. (n=7-11; KW test p-values displayed). **b)** Example EM images of myocardium from male ob/CMS mice treated with either therapy. Upper shows typical enlarged mitochondria with reduced density observed in the placebo group. (Replicated x4).

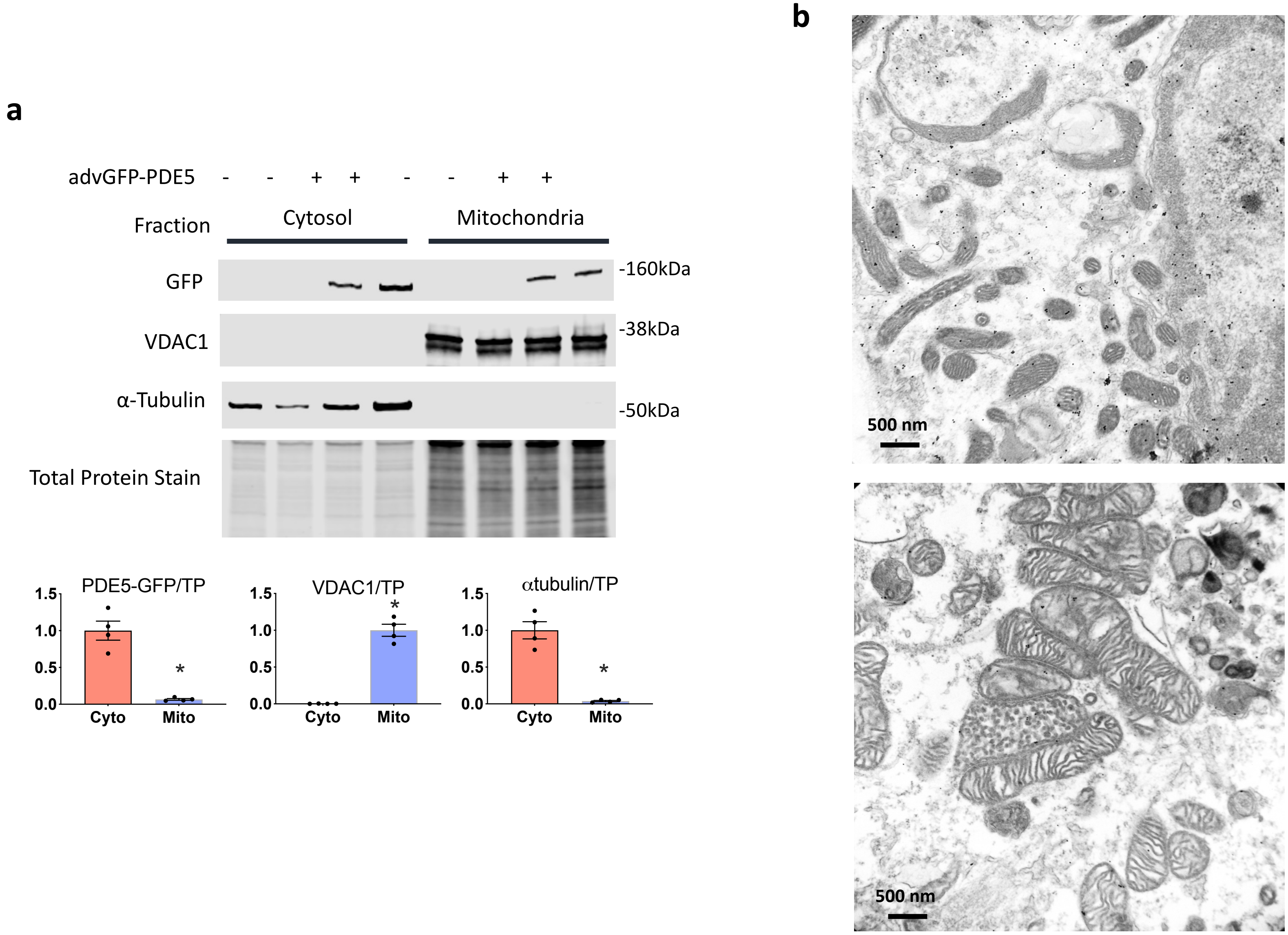
a
advGFP-PDE5 - - + + - - + +
Fraction
Cytosol
Mitochondria
GFP
-160kDa
VDAC1
-38kDa
α-Tubulin
-50kDa
Total Protein Stain
PDE5-GFP/TP
VDAC1/TP
αtubulin/TP
1.5
1.0
0.5
0.0
Cyto
Mito
*
b
500 nm
500 nm

**Figure S6**

**Figure S6. PDE5A localizes to cytosol, whereas PDE9 localizes to mitochondria. a)** Example gel and summary data for myocyte fractionation assay in cells expressing PDE5A-GFP. The fusion protein was largely constrained in the cytosolic fraction, opposite to that for PDE9 (e.g. compared with Figure 5a. (n=4, MW test, * p=0.28). **b)** EM micrographs with immunogold staining for PDE9-GFP or GFP alone. These lower power images (compare with Figure 5c) show diffuse distribution of GFP, whereas the PDE9-GFP localizes far more to mitochondria.

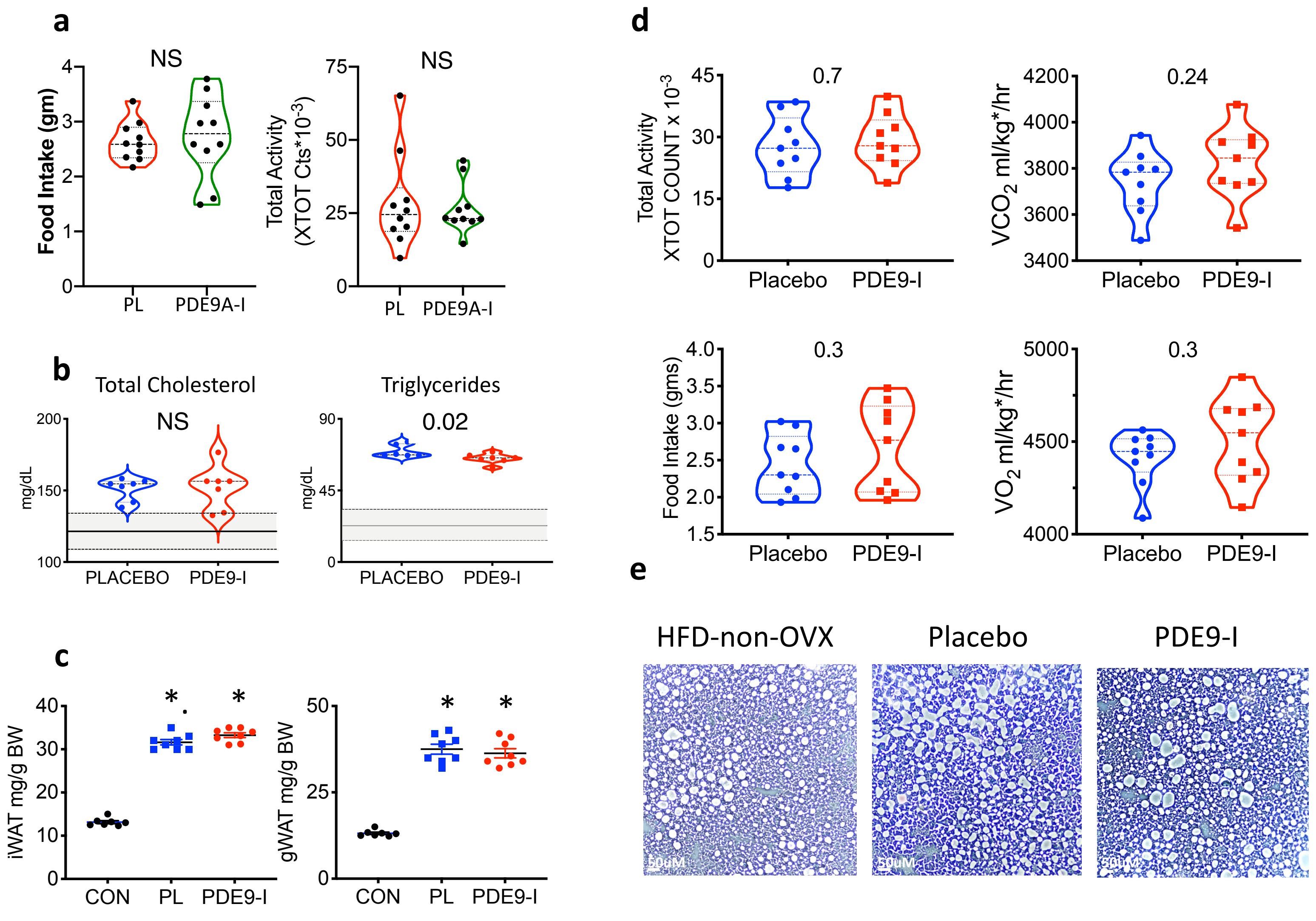
a
NS
Food Intake (gm)
PL
PDE9A-I
NS
Total Activity (XTOT Cts*10-3)
PL
PDE9A-I
b
Total Cholesterol
NS
mg/dL
PLACEBO
PDE9-I
Triglycerides
0.02
mg/dL
PLACEBO
PDE9-I
c
iWAT mg/g BW
CON
PL
PDE9-I
gWAT mg/g BW
CON
PL
PDE9-I
d
Total Activity XTOT COUNT x 10-3
0.7
Placebo
PDE9-I
VCO2 ml/kg*/hr
0.24
Placebo
PDE9-I
Food Intake (gms)
0.3
Placebo
PDE9-I
VO2 ml/kg*/hr
0.3
Placebo
PDE9-I
e
HFD-non-OVX
Placebo
PDE9-I

**Figure S7**

**Figure S7. PDE9-I +/- PPARa-I does not alter food intake or activity. Non-OVX females display no significant changes in serum lipids, iWAT or gWAT weight, food intake, activity, or whole body metabolism, or hepatic lipid accumulation in response to PDE9-I. a)** Food intake and activity OVX obese/CMS mice treated with either PDE9-I alone or in combination with PPARα-I show no significant differences in total daily activity or food intake. **b-e**) Non-OVX mice treated with placebo or PDE9-I show minimal differences in serum lipids **(b)**, iWAT or gWAT mass **(c)**, activity, food intake, $VO_2$ or $VCO_2$ (normalized to lean + 0.2x fat mass) **(d)**, or lipid accumulation in the liver **(e)**.

**Supplemental Data Table. Clinical Features of two HpPEF groups defined by relative ratio of *PPARA* and *PDE9* gene expression.**

| | Group 1: ↓ *PPARA* vs *PDE9A* (n=18) | Group 2: ↑ *PPARA* vs *PDE9A* (n=23) | P-value |
|---|---|---|---|
| Age, years | 62.5 (53.2, 66.8) | 62.0 (53, 72) | 0.48[1] |
| Female Sex, n (%) | 5 (27.8%) | 12 (52.2%) | 0.20[2] |
| African-American, n (%) | 13 (72.2%) | 15 (65.2%) | 0.64[2] |
| **NYHA Class** | | | 0.73[2] |
| II | 6 (33.3%) | 8 (34.8%) | |
| III | 11 (61.1%) | 15 (65.2%) | |
| ACEi or ARB, n (%) | 12 (66.7%) | 14 (60.9%) | 0.75[2] |
| Beta Blocker, n (%) | 9 (50.0%) | 13 (56.5%) | 0.76[2] |
| Aldosterone Blocker, n (%) | 9 (50.0%) | 4 (17.4%) | **0.04[2]** |
| Loop Diuretic, n (%) | 16 (88.9%) | 23 (100.0%) | 0.19[2] |
| **Past Medical History** | | | |
| Hypertension, n (%) | 17 (94.4%) | 23 (100.0%) | 0.44[2] |
| Diabetes, n (%) | 8 (44.4%) | 18 (78.3%) | **0.048[2]** |
| Coronary artery disease, n | 3 (16.7%) | 1 (4.3%) | 0.30[2] |
| BMI, kg/$m^2$ | 40.3 (37.5, 44) | 43.6 (34.2, 46.5) | 0.92[1] |
| SBP (mmHg) | 150.5 (124.8, 170.5) | 138.0 (133.0, 147.0) | 0.82[1] |
| DBP (mmHg) | 77.5 (72.2, 82.5) | 67.0 (64.0, 77.0) | 0.09[1] |
| LVEF, % | 65.0 (56.2, 68.8) | 65.0 (60.0, 67.5) | 0.74[1] |
| LVEDD, cm | 4.2 (3.8, 4.7) | 4.8 (4.2, 5.4) | **0.02[1]** |
| SA - LV mass/$height^{1.7}$, g/$m^{1.7}$ | 95 (86, 115) | 107 (83, 131) | |
| LVMI, g/$m^2$ | 89.4 (69.9, 102.0) | 106.1 (82.6, 130.8) | **0.05[1]** |
| NT-proBNP, pg/mL | 125.5 (48.0, 197.0) | 434.0 (104.5, 1352.0) | **0.05[1]** |
| PA Systolic Pressure, mmHg | 36.5 (32.0, 50.8) | 48.0 (41.0, 57.5) | **0.04[1]** |
| **Invasive Hemodynamics** | | | |
| PAWP, mmHg | 20.0 (14.2, 23.5) | 20.0 (15.5, 24.0) | 0.72[1] |
| PVR (mmHg/L/min) | 1.3 (0.9, 1.8) | 2.0 (1.3, 2.7) | **0.02[1]** |
| CO, L/min | 6.0 (5.1, 6.6) | 5.9(4.8, 7.0) | 0.92[1] |
| **NMF: Sub-Group** | **NMF: Group 1 (LVH,** | **NMF: Group 2 (Small** | |
| Group 1 (↓*PPARA/PDE9*) | 0 (0%) | 8 (53%) | |
| Group 2 (↑*PPARA/PDE9*) | 11 (48%) | 2 (8.7%) | **0.0005** |

Data are n (%) or median (25th-75th percentile). P-value displayed for Fisher's exact test used for categorical variables, otherwise Mann Whitney comparison test. ACEi, angiotensin converting enzyme inhibitor; ARB, angiotensin II receptor blocker. HF, heart failure; NYHA, New York Heart Association; ACEi, angiotensin converting enzyme inhibitor; ARB, angiotensin II receptor blocker; BMI, body mass index; LVEF, left ventricular ejection fraction; LVEDD, left ventricular end diastolic diameter; LV, left ventricle; SA – sex adjusted LV mass/height$^{1.7}$ calculated by multiplying by a constant of 1.28 for women; eGFR, estimated glomerular filtration rate; NTproBNP, N-terminal pro-B type natriuretic peptide; RAP, right atrial pressure; Mean PAP, mean pulmonary artery pressure; PAWP, pulmonary artery wedge pressure; CO, cardiac output; CI, cardiac index; SVRI – systemic vascular resistance index.
